# TIRR regulates mRNA export and association with P bodies in response to DNA damage

**DOI:** 10.1101/2024.02.19.580988

**Authors:** Michelle S Glossop, Irina Chelysheva, Ruth F Ketley, Adele Alagia, Monika Gullerova

## Abstract

To ensure the integrity of our genetic code, a coordinated network of signalling and repair proteins known as the DNA damage response (DDR) detects and repairs DNA insults, the most toxic being double-stranded breaks (DSBs). Tudor interacting repair regulator (TIRR) is a key factor in DSB repair, acting through its interaction with p53 binding protein 1 (53BP1). TIRR is also an RNA-binding protein, yet its role in RNA regulation during the DNA damage response remains elusive. Here we show that TIRR selectively binds to a subset of mRNAs in response to DNA damage with preference for transcripts encoding transcription factors and RNA polymerase II (RNAPII) transcription regulators. Upon DNA damage, TIRR interacts with the nuclear export protein Exportin-1 (XPO1), through its nuclear export sequence (NES). Furthermore, TIRR plays a crucial role in modulation of RNA processing bodies (P bodies/PBs). TIRR itself and TIRR-bound RNA co-localises with PBs, and TIRR depletion results in nuclear RNA retention and impaired PB formation. Finally, the role of TIRR in RNA export contributes to efficient DNA damage response. This work reveals intricate involvement of TIRR in orchestrating mRNA nuclear export and storage within PBs, emphasizing its significance in the regulation of RNA-mediated DNA damage response.

## Introduction

A large network of signalling and repair proteins known as the DNA damage response (DDR), acts to ensure efficient repair caused by various damaging insults, including those resulting in DNA double-stranded breaks (DSBs). Among these proteins, TIRR, a NUDIX family protein (1), sits at the interface between homologous recombination (HR) and non-homologous end joining (NHEJ), two key pathways in DSB repair (2). TIRR binds and inhibits 53BP1, an integral factor in promoting NHEJ (3,4). 53BP1 counteracts resection, a step which is required to initiate HR, and is therefore extensively regulated. The role of TIRR in 53BP1 binding and regulation in the DSB repair response has been well characterised (1,5–9). In non-damage conditions, TIRR and 53BP1 form a complex. In response to DSBs, TIRR and 53BP1 dissociate, allowing 53BP1 to bind chromatin at the break (5). However, the wider role of TIRR outside of 53BP1 binding has not been extensively explored, especially in the context of DNA damage.

TIRR has been characterised as an RNA binding protein (6–8), binding a wide range of RNA, particularly mRNA (6). Initial characterisation of the role of TIRR in RNA binding in non-damage conditions suggests that TIRR can act as a translational repressor of the RNA to which it binds (6). Interestingly, TIRR was also identified as a pre-miRNA binding protein in *in vitro* capture and mass spectrometry experiments, binding multiple different precursor microRNA (pre-miRNA) (9). Additionally, TIRR RNA binding is relevant in the context of DNA damage. We have previously shown that hairpin shaped RNA derived from the DSBs contributes to the dissociation of TIRR from 53BP1 after DSB induction (8). Therefore, TIRR binds to a broad spectrum of RNA with distinct functional outcomes, yet this has been studied limitedly in the context of DNA damage.

RNA regulation and processing, including RNA transcription, stability, translation, storage, and degradation can be defined with the general term of RNA metabolism(10). RNA metabolism is extensively regulated in DDR, including global transcription inhibition of mRNA, alterations in splicing and degradation of transcripts, and production of damage-specific RNA species (11–15). Many different proteins act to regulate RNA metabolism, carefully coordinating RNA fate in different cellular contexts. For example, NUDIX family proteins participate in RNA metabolism (1), one of which being NUDT16, a close paralogue of TIRR involved in RNA de- capping (16). The regulation of RNA fate, such as decay and storage, can occur in different cellular compartments, including Processing bodies (P-bodies or PBs). PBs are cytoplasmic membrane-less compartments which contain RNAs and various RNA-binding proteins (RBPs). In general, PBs are considered to be sites of non-translating mRNA(17) and sites of RNA decay or storage(17–19), promoting cell survival during and after stress(20,21). PBs contain components of the mRNA degradation machinery, such as DCP1/2, XNR1, and the CCR4- NOT complex, which are involved in mRNA de-capping and de-adenylation, as well as components of the miRNA pathway, such as GW182 and miRNAs(22). PBs can also associate with polysomes(19) and stress granules (SGs) (23), highly dynamic sites of mRNA storage and concentrated translation initiation factors. mRNAs can move between polysomes, where they are translated, and PBs, where they can be either stored or degraded, and SGs where they may be reintroduced to translation initiation factors, in a model known as the mRNA cycle (19,21,24).

Given that TIRR is an RNA binding protein involved in DSB repair, and DNA damage is known to impact RNA metabolism/functions, we wanted to assess whether TIRR may play a role in regulation of RNA metabolism in response to DNA damage. As little is known about the function of TIRR in RNA regulation. TIRR could potentially participate at any stage of RNA metabolism, such as transcription, splicing and export from the nucleus, translation into proteins, and RNA decay(20). To investigate this, we first performed RNA immunoprecipitation and sequencing (RIP-Seq) experiments to identify specific RNAs bound to TIRR after DNA damage. Our analysis revealed that TIRR selectively binds to a subset of mRNAs involved in the regulation of RNAPII transcription. Furthermore, we demonstrate that a fraction of TIRR is exported from the nucleus through XPO1 binding upon DNA damage. TIRR localises to P bodies with its bound mRNAs and participates in the regulation of their formation in response DNA damage.

In essence, TIRR emerges as a pivotal player orchestrating the intricate features of mRNA regulation in the wake of genomic insults.

## Materials and Methods

### Cell Culture

Flp-IN TIRR-GFP, GFP, shTIRR, shGFP cells, Wild type HeLa cells and Wild type U2OS cells were cultured in DMEM containing 1% L-glutamine, and 10% FBS at 37°C with 5% CO_2_ with and without antibiotics. TIRR-GFP, GFP, shTIRR, and shGFP cells were a gift from Rosario Avolio. Drugs were added to cell culture media as indicated: Doxycycline (1μg/ml, 16-48 hours), Etoposide (ETO) (5-10μM, 2 hours), α-amanitin (2μg/ml, 24 hours), and Leptomycin B (LMB) (5nM, 24 hours). Double strand breaks were induced using ionising radiation (IR) at 10 Gy using a Gravatom (GRAVITRON RX 30/55), followed by incubation at 37°C for the specified times.

### Cloning and Plasmids

All cloning was performed using Gibson cloning with the NEBuilder® HiFi DNA Assembly Master Mix (New England Biolabs). The PAGFP-TIRR plasmid was generated by amplifying a PAGFP fragment from a pPAGFP-C1 plasmid and a backbone containing TIRR using Q5® High-Fidelity DNA Polymerase and associated Q5® Reaction Buffer and GC enhancer (New England Biolabs). The PAGFP fragment was inserted C-terminally to TIRR.

pPAGFP-C1 was a gift from Jennifer Lippincott-Schwartz (Addgene plasmid # 11910 ; http://n2t.net/addgene:11910; RRID:Addgene_11910) (25). The LSM14A-mScarlet plasmid was generated similarly, inserting mScarlet from an existing TIRR-mScarlet plasmid N-terminally into a LSM14A-containing backbone derived from an LSM14A-GFP plasmid. The LSM14A-GFP plasmid was a gift from the Weil Lab at Sorbonne Université (24).

Due to the GC rich sequence of TIRR, the RNA-binding mutant and NES mutants were generated by another method. For the RNA-binding mutant, forward and reverse 180nt ssDNA fragments with overlapping 3’ complementary regions were designed to contain individually mutated residues according to (26). For the NES mutants, each ssDNA fragment contained either a wildtype TIRR sequence or a mutated NES site, in which all hydrophobic residues were mutated to alanine (alanine-scanning) and the surrounding TIRR sequence. All ssDNA fragments were produced as Ultramers™ from IDT. These ssDNA fragments were then annealed and extended to form a double-stranded DNA fragment containing either RBM TIRR or mutated NES sites that were sufficiently large for efficient Gibson cloning. These fragments were amplified along with a GFP-containing plasmid backbone. Cloning was performed as above. mCherry-LaminB1-10 was a gift from Michael Davidson (Addgene plasmid # 55069 ; http://n2t.net/addgene:55069; RRID:Addgene_55069). For oligonucleotide sequences see Supplementary Table 3.

### Western Blot

Western blotting was performed as in (8). Protein samples were boiled in Laemelli buffer for 7 minutes at 98°C, run on 4-15% Mini-Protean TGX gels (BioRad) at 100 V for approximately 1.5 hrs and transferred onto nitrocellulose membranes using a Trans-Blot Turbo Transfer System (BioRad). Membranes were blocked in 5% Milk in PBST for 1 hour, before incubation with primary antibodies overnight at 4°C. Secondary antibody was incubated for 1 hour at room temperature (RT). Membranes were visualised using ECL (Thermo) and developed on Amersham Hyperfilm ECL film. Primary antibodies used: GFP Monoclonal antibody (3H9, Chromotek) 1:1000-2000, Rb pAb to beta-tubulin (Abcam) 1:2000-4000, GAPDH Monoclonal antibody (Proteintech) 1:2000-4000, Rabbit anti-53BP1 pAb (Novus Biologicals) 1:500, Anti-phospho-Histone H2A.X (Ser139) Antibody clone JBW301 (Millipore) 1:1000, Anti- NUDT16L1 (Atlas Antibodies) 1:500, Anti-LSM14A Purified MaxPab antibody (Novus Biologicals) 1:250, CRM1 C-1 antibody (Santa Cruz) 1:250, mCherry antibody 1:1000 (GeneTex), Rb pAb to LaminB1 (Abcam) 1:1000. NUDT16L1 antibody (Atlas Antibodies) and CRM1 antibody (Santa Cruz) were also incubated with SuperSignal™ Western Blot Enhancer (Thermo Fisher).

### RNA immunoprecipitation and Sequencing (RIP-Seq)

3 x 15-cm dishes were used per condition and plated to achieve ∼70% confluency at the point of harvest. TIRR-GFP and GFP cells were plated and 24 hours later, induced with 1 μg/ml doxycycline overnight to induce expression. Cells were washed three times in cold PBS and cross-linked at 254nm at 0.15 J/cm^2^. Cells were lysed in cold RIP lysis buffer (100mM NaCl, 15mM MgCl_2_, 10mM Tris pH 7.5, 0.5% NP40, 0.5% Sodium Deoxycholate, 1mM DTT) with 1 x protease inhibitor (Roche) and Ribolock (Thermo), with Turbo DNase (Invitrogen). Lysates were sonicated (10s on, 30s off) at medium intensity. Lysates were centrifuged at 17,000g for 10 minutes and the supernatant collected. GFP-Trap A beads were washed 3 times in lysis buffer and added to samples for 2 hours on a wheel at 4°C. Beads were collected by centrifugation at 2500g for 2 minutes and washed 2x with RT RIP high salt buffer (500mM NaCl, 1mM MgCl_2_, 20mM Tris pH 7.5, 0.05% NP40, 0.1% SDS, Protease inhibitor), 2x with cold RIP low salt buffer (150mM NaCl, 20mM Tris pH 7.5, 1mM MgCl_2_, 0.01% NP40), and 2x with cold RIP Proteinase K buffer (100mM Tris pH 7.5, 50mM NaCl, 10mM EDTA). Beads were incubated with Proteinase K (Thermo) in RIP Proteinase K buffer at 37℃ for 1 hour with shaking at 1100rpm. RNA was extracted using Trizol and the Monarch Total RNA Miniprep kit according to the manufacturer’s instructions. RNA samples were submitted to the Oxford Genomics Centre for rRNA depletion and paired-end bulk full transcript RNA sequencing on the Illumina NovaSeq6000 was performed.

### Total RNA extraction

Total RNA was extracted by scraping from plates using RNA lysis reagent from the Monarch Total RNA Miniprep Kit. RNA was eluted in water and quantified using an Implen NanoPhotometer N60.

### RNA-sequencing

Total RNA-sequencing was performed on stably integrated inducible shRNA HeLa cells, expressing shRNA for TIRR or GFP. TIRR knockdown was induced using doxycycline for 48 hours. RNA was collected before and after DNA damage, induced by IR and followed by 1 hour incubation, or by ETO incubation for 2 hours. It was then subjected to paired-end total RNA sequencing in three replicates at Azenta Life Sciences on the Illumina NovaSeq6000.

### Fluorescence *in situ* hybridisation (FISH) FISH/IF

TRex Flp-IN shTIRR and TRex Flp-IN shGFP HeLa cells were plated on poly-L-lysine coated glass cover slips. TIRR knockdown was obtained by a doxycycline-inducible short hairpin RNA (shRNA) against TIRR (Avolio et al., 2018). Coverslips were then washed 3 times in cold PBS and fixed in cold 4% paraformaldehyde (Alfa Aesar) in PBS for 15 min. Coverslips were then washed 3 times in cold PBS and permeabilised with cold 0.2% Triton X-100 in PBS (PBST) for 10 min. Slides were then washed 3 times in cold PBS and blocked in FISH blocking buffer (2x SSC, 40ng/μL yeast tRNA (Life Technologies), 40ng/μL salmon sperm DNA (Life Technologies), 0.5% Triton X-100, 2% BSA, 0.7U/μL RNAsin Plus (Promega)) at 37°C for 1 h. Coverslips were incubated with FISH probe incubation buffer (2x SSC, 40ng/μL yeast tRNA (Life Technologies), 40ng/μL salmon sperm DNA (Life Technologies), 0.5% Triton X-100, 2% BSA, 1U/μL RNAsin Plus (Promega) and 100nM DNA probes) for 3 min at 95°C, and then at 4°C overnight in a humidified chamber. Coverslips were washed 3 times for 10 minutes at room temperature in 2x SSC-T buffer (2x SSC, 0.05% Triton X-100), 3x in 1x SSC-T buffer (1x SSC, 0.05% Triton X-100), 3x in 0.5x SSC-T buffer (0.5x SSC, 0.05% Triton X-100) and 3x in 0.25x SSC-T buffer (0.25x SSC, 0.05% Triton X-100). Coverslips were then incubated with IF blocking buffer (2xSSC, FBS 5%, BSA 5%) for 1 hour at 37°C. Primary antibodies (LSM14A (18336-1-AP, Proteintech) and G3BP1 (13057-2-AP, Proteintech)) were diluted to the appropriate concentration in 5% FBS and 5% BSA in 2xSSC and incubated overnight at 4°C in a humidified chamber. Coverslips were then washed 3 times in SSC-T buffer (2xSSC, 0.1% Triton X-100) and subsequently incubated for 2 h at RT with Donkey anti-Rabbit Alexa Fluor™ 555 (A32794, Invitrogen) secondary antibodies diluted in 0.1% FBS and 0.1% BSA in 2x SSC. Coverslips were washed 3 times in 2x SSC 0.1% Triton X-100 and once in 2x SSC before mounting on slides with Fluoroshield Mounting Medium with DAPI (Abcam).

DNA FISH Probes

Peli2: Alexa Fluor 488 – AAGCACCATTGTACCCGAGC

ZFN600: Alexa Fluor 488 – TGCGTTTGGAAGAGATATCCAC

SPEN: Alexa Fluor 488 – AGCGCTCCAAACGAGACTTG

Tubulin: Alexa Fluor 488 – CAGAGTCCATGGTCCCAGGT

### Confocal Microscopy

Immunofluorescence and confocal microscopy were performed according to a previously published protocol(8). Cells were plated on glass coverslips and fixed in cold 4% paraformaldehyde in PBS for 10 minutes. Cells were permeabilised with cold 0.2% Triton X- 100 in PBS for 10 minutes, followed by blocking in 10% FBS in PBS for 2 hours at 4°C. Primary antibodies were diluted in 10% FBS in PBS and incubated overnight at 4°C in a humidified chamber. Secondary antibody incubation was performed for 2 hours at RT with secondary antibodies diluted in 0.15% FBS in PBS. Fluoroshield Mounting Medium with DAPI was used to mount slides on coverslips, and imaging performed using an Olympus Fluoview FV1200 confocal microscope with a 60X objective lens.

### Photoactivation and Live Cell Microscopy

Live cell imaging was performed with a SoRa spinning disc confocal microscope. U2OS cells were reverse transfected with 1 ug each of PAGFP-TIRR and mCherry-LaminB1-10 or LSM14A-mCherry plasmids of Lipofectamine LTX with PLUS reagent (ThermoFisher) in plastic 6-well plates and then reseeded on poly-L-lysine coated glass bottom 35mm plates. Cells were treated with LMB where stated, and damage was induced the following day with IR. Plates were immediately brought to the microscope for imaging. PAGFP-TIRR was induced by 405 nm laser, which was followed by time-lapse imaging, using 488 nm and 555 nm lasers for PAGFP-TIRR and mCherry-LaminB1-10 respectively. Images were taken every 4 sec for 2 minutes. Quantification of the cytoplasmic signal was performed using FIJI software.

### Proximity Ligation Assay (PLA)

Proximity Ligation Assay was performed according to the manufacturer’s instructions and our previously published protocol(8).

### yH2AX immunofluorescence time course

Approximately 800,000 U2OS cells were reverse transfected with 20 nM siRNA against the 3’ UTR of endogenous TIRR or a negative control siRNA (Dharmacon ON-TARGETplus Human NUDT16L1 siRNA J-014872-23-0005 or ON-TARGETplus Non-targeting Control) using 3 uL of RNAiMAX lipofectamine (Thermo Fisher) in a 6-well plate. After 24 hours, the cells were split into two wells and reverse transfected with 1 ug of plasmid each, corresponding to a GFP only control, TIRR-WT-GFP, TIRR-RBM-GFP, TIRR-K10E-mNeonGreen (4), and TIRR-NES2-GFP. 5 uL of Lipofectamine LTX was used with 1 uL of PLUS reagent for transfection. 24 hours later, the cells were split into both 35 mm plates and poly-lysine coated coverslips in 35mm plates for western blot and immunofluorescence analysis, corresponding to 3 plates per condition for three time points. 12 hours later, cells were irradiated with 10 Gy IR. Samples were collected at 15 minutes, 6 hours and 24 hours and processed according to the appropriate methods. A BCA assay (Thermo) was used to evaluate the protein concentrations of samples and concentrations were normalized accordingly.

### Co-immunoprecipitation (co-IP)

Co-IP was performed according to a previously published method (8). For NES mutant analysis, approximately 2 million U2OS cells were reverse transfected with 2 ug of the corresponding plasmid (GFP only, TIRR WT, TIRR RBM, and NES1, NES2, NES3 or NES4) using Lipofectamine LTX with PLUS reagent (ThermoFisher) in 6-well plates and then reseeded on 15 cm dishes the following day. Briefly, cells were lysed in lysis buffer (50mM Tris pH 7.5, 150mM NaCl, 1mM EDTA, 5mM MgCl_2_, 0.5% NP40, Protease and Phosphatase inhibitors (Roche), Pierce Universal Nuclease (Thermo)) on a wheel at 4°C for 30 minutes, before centrifugation at 17,000g for 10 minutes. Lysates were diluted in lysis buffer without NP40 (1.5X cell lysate volume), and incubated with GFP-Trap magnetic beads (Chromotek) for 1.5 hours at 4°C. Beads were washed in lysis buffer without NP40 and samples were eluted using 2x Laemmli buffer and boiling at 95°C.

### In vitro binding assay of TIRR and XPO1

Ni-NTA Magnetic Agarose Beads were incubated with 1.5µg of recombinant His-TIRR(8) in binding buffer (50mM Tris pH 7.4, 150mM NaCl, 5mM MgCl_2_, 10% Glycerol, Protease inhibitors (EDTA free, Roche)) for 1 hour at 4°C on a wheel. Beads were washed 2x in binding buffer, and incubated with 6µg recombinant GST-tagged XPO1 (ab131897, Abcam) for 1 hour at 4°C on a wheel. The supernatant was removed and the beads washed 4x in binding buffer. Proteins were eluted in 500mM imidazole and eluant was collected and ran on a western blot to visualise binding.

### Details about oligonucleotides used in this study can be found in Supplementary Table 3

#### Data and statistical analysis

##### RNA Sequencing analysis

The sequencing data was aligned against the whole human (*Homo sapiens*) genome build GRCh38 (https://ccb.jhu.edu/software/hisat2/index.shtml), using STAR (version 2.7.3a). Gene features were counted using HTSeq (version 0.11.1), using human gene annotation general transfer format version GRCh38.92 (www.ensembl.org). Genes with low counts across most libraries were removed. Ribosomal RNA (rRNA), sex chromosome genes, mitochondrial RNA and haemoglobin genes were excluded from downstream analysis. Differential gene expression was conducted using the R Bioconductor packages “edgeR” and “limma”(27–30). RNA-sequencing data were then normalised for RNA composition using the trimmed mean of M-value (TMM) method(29). Data were transformed using the limma “voom” function. A linear model was fitted to the data using the limma “lmFit” function using the empirical Bayes method(28). The cut-off for statistical significance was set at false-discovery rate (FDR) <0.05. PCA analysis and heatmaps were performed using the R Bioconductor package “pcaExplorer”(31) on the log transformed data; confidence interval level of 0.95 has been used. GO-terms enrichment analysis and characterisation of the gene lists compared to the whole genome were performed in ShinyGO (version 0.76)(32) using GO Biological process and GO Molecular function databases, where specified. The cut-off for statistical significance was set at false-discovery rate (FDR) <0.05, up to 20 top pathways were shown with redundant pathways removed.

### Statistical Tests

To test for a normal distribution, the Kolmogorov-Smirnov normality test was performed. If data met the requirements of a normal distribution, an unpaired t-test was performed. For comparison of two groups if data did not show a normal distribution, a Mann-Whitney test was performed. For comparison of more than two groups, a Kruskal-Wallis with Dunn’s multiple comparisons test was carried out. P>0.05 = ns, p≤0.05 = *, p≤0.01 = **, p≤0.001=***, p≤0.0001=****. GraphPad Prism 9 was used to perform the statistical analysis.

## Results

### TIRR binds to transcripts of proteins associated with RNA polymerase II transcription regulation in response to DNA damage

To explore how TIRR may function as an RNA-binding protein in DNA damage, we first performed RNA immunoprecipitation with UV crosslinking followed by sequencing (RIP-Seq), allowing us to identify specific RNA bound to TIRR in DNA damage conditions (33–35) (**Supplementary Figure S1A)**. UV light at 254 nm directly crosslinks RNA and protein, but not protein and protein, allowing for stringent washing conditions during immunoprecipitation to remove nonspecific RNA. To perform the RIP-seq, TIRR-GFP or GFP only (to control for background) was overexpressed through doxycycline induction in HeLa cells with integrated inducible TIRR-GFP or GFP only constructs (referred to as TIRR-GFP OE and GFP OE cell lines) (**Supplementary Figure S1B**). This was followed by the introduction of DSBs using etoposide (ETO), and subsequent RIP-Seq (**Supplementary Figures S1C-G).** Principal Component Analysis (PCA) revealed good reproducibility of the RIP-seq samples, although there is some variability in the samples isolated from TIRR-GFP OE non-damage replicates (**Supplementary Figure S1F**). However, utilising the Euclidean distance on the heatmap, the samples from the same condition do cluster together (**Supplementary Figure S1D**). The potential variability observed in TIRR non-damage samples may reduce the number of significantly differently expressed genes for this condition compared to the GFP non-damage condition, but more importantly will not affect the comparisons performed in the damaged condition and targets detected in TIRR damage. In total, we identified 729 RNAs bound to TIRR in non-damage conditions, and 1207 RNAs bound to TIRR after damage. Intersection of these RNAs revealed an overlap of 572 RNAs. Interestingly, we identified 635 RNAs uniquely bound to TIRR upon DNA damage (**Figure 1A-C and Supplementary Table 1**). GO-term analysis of TIRR bound RNAs found both in damage and non-damage conditions showed an enrichment in biological processes including mRNA processing and mRNA splicing, suggesting a potential role for TIRR in RNA metabolism (**Supplementary Figure S2A**). GO analysis of RNAs which were bound to TIRR specifically after damage showed an enrichment for pathways involved in the regulation of RNAPII transcription, including: DNA-binding transcription factor activity, sequence specific DNA binding, and RNAPII transcription regulatory region sequence specific binding (**Figure 1D and Supplementary Figure S2B**). Interestingly, among these mRNA, we identified a subset of zinc finger containing transcription factors, predominantly encoded on chromosome 19 (**Figure 1E and Supplementary Figure S2C**). Further analysis of RNAs bound to TIRR after damage showed no significant difference in 5’UTR length, transcript length, or 3’UTR length relative to the rest of the genome (**Supplementary Figure S3**). However, analysis of the GC content of RNAs bound to TIRR revealed a significant enrichment of mRNAs with lower GC content (**Figure 1F),** which could be relevant to their metabolism (18).

**Figure 1:**
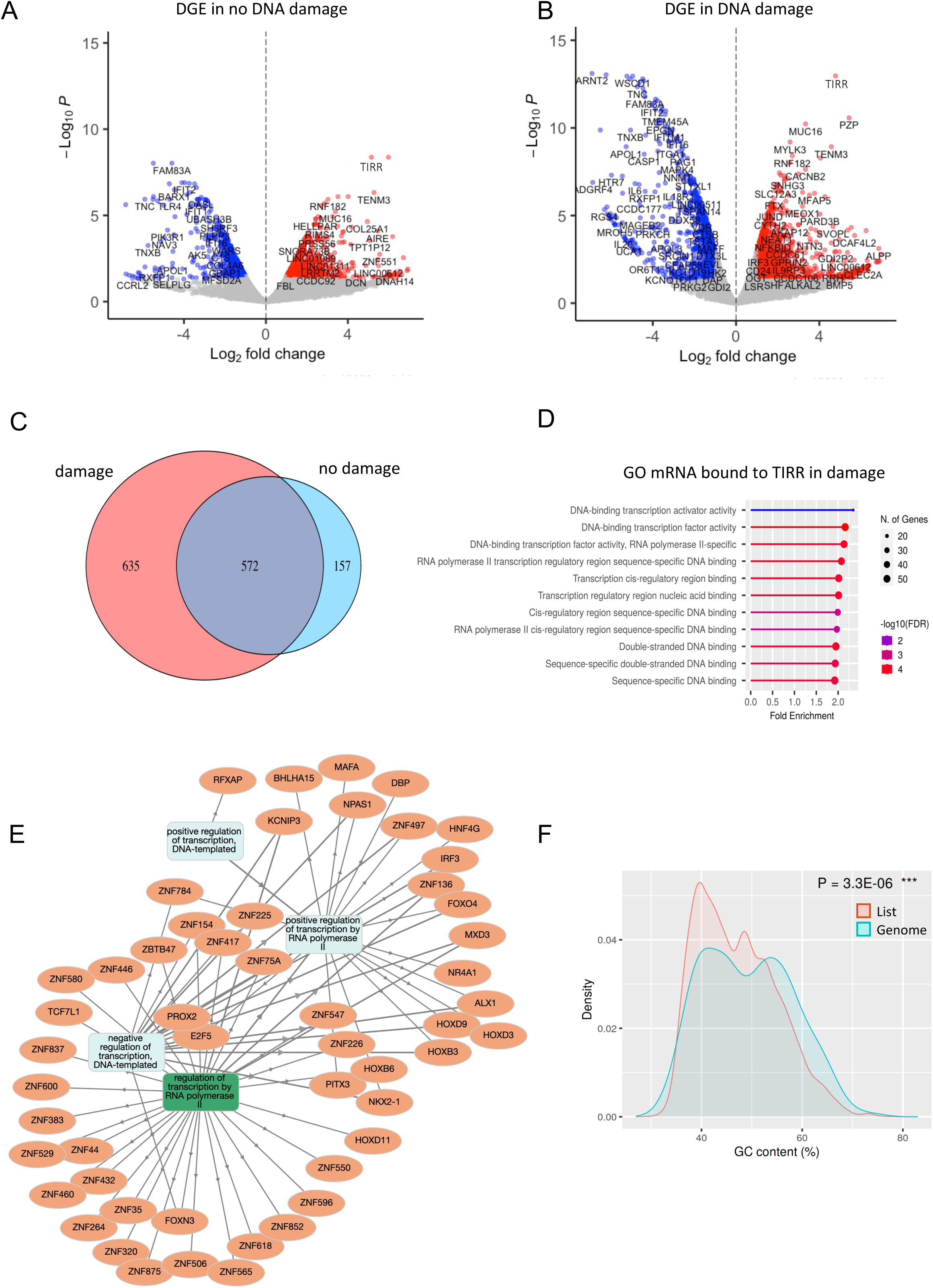
TIRR binds to mRNA of transcription associated proteins upon DNA Damage. A) Volcano plot showing differentially enriched RNAs in TIRR-GFP IP over GFP only IP in non-damaged conditions. TIRR-GFP IP (n=2), GFP IP (n=1). B) As in A, in DNA damage conditions. C) Pie chart showing an overlap between RNA transcripts identified by TIRR RIP-seq in non- damage conditions and DNA damage conditions. D) GO analysis of RNAs bound to TIRR in DNA damage conditions. E) Network presentation of RNAs bound to TIRR in damage conditions, which were identified through GO analysis as significantly enriched in RNAPII transcription regulation. F) Comparison of GC content feature of RNAs bound to TIRR in damage conditions vs the reference genome.

To assess what role TIRR may serve in binding to this subset of mRNA corresponding to RNAPII regulatory factors, we first performed total RNA-Seq in stably integrated doxycycline-inducible shRNA HeLa cells, expressing shRNA for TIRR (shTIRR) or GFP (shGFP) as a control. TIRR knockdown was induced using doxycycline for 48 hours, before inducing damage by treatment with ETO (5 uM) for 2 hours. Total RNA was collected and subjected to paired-end total RNA sequencing. Heatmap analysis and PCA revealed similarity between the RNA-seq replicates (**Supplementary Figure S4A and B).** We observed that TIRR RNA was successfully depleted in our sequencing experiments, due to a significant decrease in the relative expression of TIRR/NUDT16L1 RNA after TIRR KD (**Supplementary Figure S4C)**. Globally, TIRR knockdown did not result in many significant changes in total RNA levels with very few RNAs showing a log_2_FC greater than 1 or less than -1 and FDR <0.05, suggesting TIRR does not have a large influence on steady-state RNA levels. Many of the same RNAs were similarly differentially expressed in TIRR knockdown in non-damage conditions compared to damage conditions (**Supplementary Figure S4D-G and Supplementary Table 2)**, suggesting TIRR knockdown neither dramatically impacts steady state RNA levels generally nor in a damage specific manner. We also performed RNA-Seq of samples in which damage was induced by IR (10Gy) followed by a 1-hour incubation period, to see if a more temporally concentrated dose of DNA damage could yield different results, however we observed a similar lack of a significant change in steady state RNA levels (**Supplementary Figure S5A-F)**. Comparison of both RNA-seq data sets (ETO and IR) revealed similarity in DEG under these two conditions (**Supplementary Figure S5G).** Comparison of the RIP-Seq data with our RNA-Seq data showed that of the 1207 RNA transcripts bound in damage conditions, only 3 were found to be significantly differentially expressed in TIRR knockdown in ETO damage compared to non-damage conditions (>1%). Overall, we conclude that TIRR binding to RNA does not dramatically influence the total steady-state levels of RNA globally. To further explore the impact TIRR binding may have on these transcripts, we also selected two proteins encoded by these transcripts to analyze by western blot in damage and TIRR knockdown conditions. HeLa cells expressing shTIRR or shGFP as a control were induced with doxycycline and treated with ETO to induce DNA damage or left untreated. In all conditions, protein levels did not change significantly (**Supplementary Figure S6A-C)**, suggesting that TIRR does not directly regulate protein production of bound transcripts.

### TIRR depletion leads to mRNA retention in the nucleus

As TIRR did not appear to influence steady-state RNA levels and selected protein levels of mRNA bound to TIRR upon damage, we instead explored whether it may impact localization of bound mRNAs and potentially influence their fate. We performed fluorescence *in situ* hybridization (FISH) using DNA probes for selected mRNAs, *ZNF600* and *PELI2*, identified to be specifically bound to TIRR in damage conditions. A DNA probe specific for *SPEN* and another one specific for *Tubulin*, both mRNAs not found to bind TIRR, were used as negative controls. FISH was performed in shTIRR and shGFP cells in non-damage and ETO-induced damage conditions. We observed that upon damage, both *ZNF600* and *PELI2* mRNAs were predominantly localised in the cytoplasm in shGFP control cells. However, depletion of TIRR resulted in significant *ZNF600* and *PELI2* mRNAs nuclear retention (**Figure 2A and B**). Such a TIRR-dependant change in localization was not observed in FISH of our negative controls, *SPEN* or *Tubulin* (**Figure 2C and D**). In the no damage conditions, nuclear retention was also observed upon TIRR knockdown, however this was less pronounced than in damage conditions (**Supplementary Figure S7A-D).** Similarly, to these observations, we also detected TIRR dependent cytoplasmic localisation of *ZNF600* and *PELI2* mRNAs upon IR treatment (**Supplementary Figure S8A and B).** These data suggest that TIRR might be required for nuclear export of specific mRNAs that it binds upon DNA damage induction.

**Figure 2:**
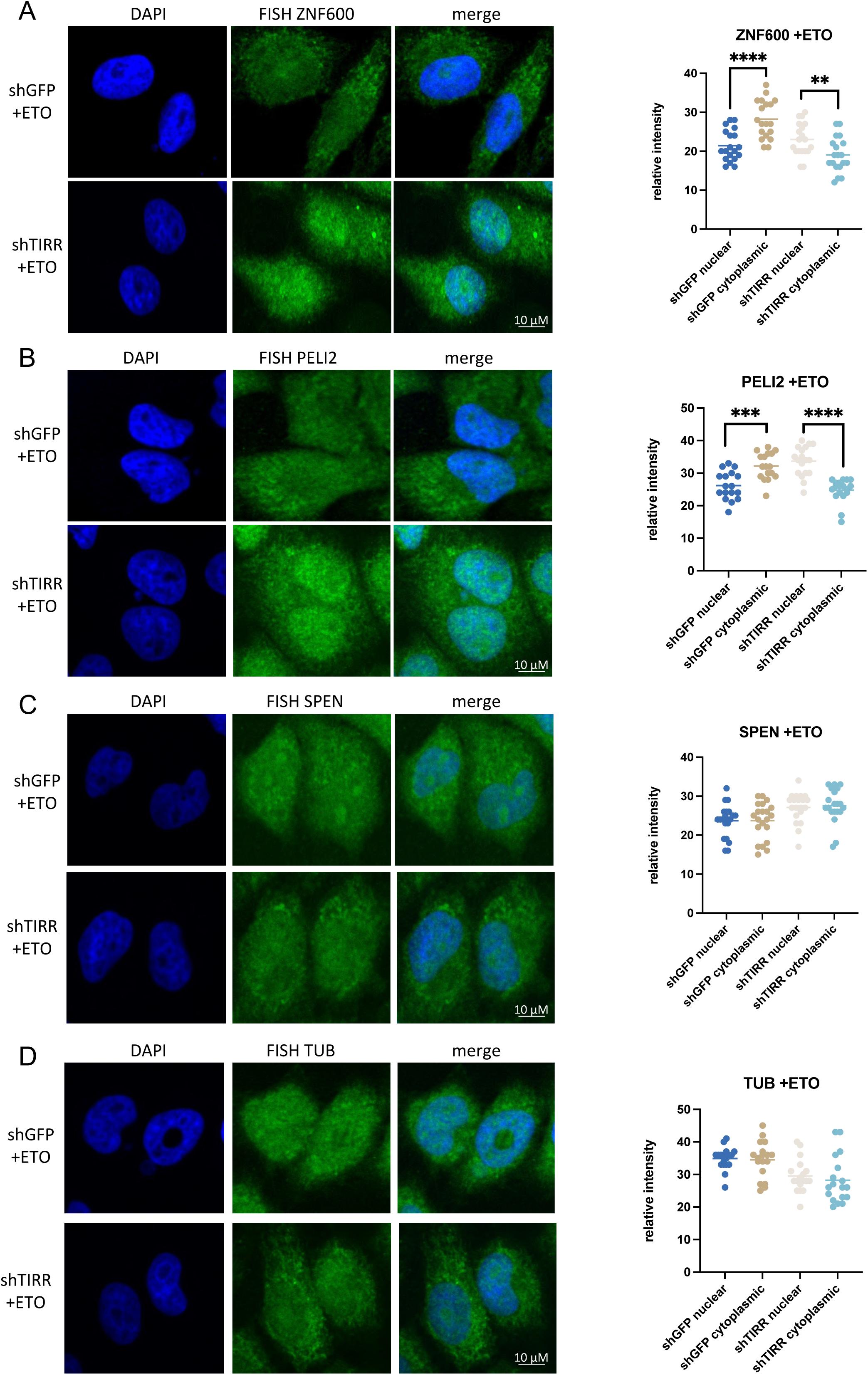
TIRR depletion leads to mRNA retention in the nucleus. A) Left: Confocal images showing RNA FISH signals (in green) for *ZNF600* in shGFP or shTIRR cells after ETO treatment. Right: Quantification of *ZNF600* mRNA signals in nucleus and cytoplasm plotted as relative FISH signal intensity in shGFP or shTIRR cells upon ETO, n>50 cells. Significance was determined using t- test, (***p ≤ 0.001, ****p ≤ 0.0001). B) As in A) for *PELI2* mRNA. C) As in A) for *SPEN* mRNA D) As in A) for *Tubulin* mRNA

### TIRR binds XPO1 upon DNA damage

Next, we tested whether there is indeed export of TIRR/mRNA complexes upon DNA damage. TIRR has been shown to exist in both the nuclear and cytoplasmic cellular compartments (5,6). Therefore, we endeavoured to determine if there were any features on TIRR relevant to its nuclear export. To address this question, we used several programmes designed to identify both general short amino acid motifs recognized by protein interactors [Eukaryotic Linear Motif (ELM) and Wregex] and specifically nuclear export signals (NES) (NESmapper and LocNES) (36–39). NES motifs can be bound by the protein chromosome region maintenance protein 1, also known as Exportin-1 (XPO1/CRM1). Exportin-1 (XPO1) is the main protein export receptor in human cells (40), regulating the export of different proteins and some RNAs, including snRNA, tRNA, and a subset of mRNAs, from the nucleus through the nuclear pore complex (NPC) into the cytoplasm (40,41). XPO1 associates with RAN-GTP to bind its cargo, exports through the NPC and, upon RAN-GTP hydrolysis to RAN-GDP, releases its cargo into the cytoplasm (41). XPO1-dependent RNA export is important for many cellular processes, such as translation and mRNA decay (42). To mediate RNA export, XPO1 does not bind directly to RNAs, but instead binds to proteins with a nuclear export signal (NES) that can themselves bind RNA, such as HuR (40,41), making XPO1 an interesting candidate for export of TIRR and its bound mRNA. Combined, ELM, Wregex, NESmapper, and LocNES identified four candidate NES motifs in TIRR (**Figure 3A**). The positions of these candidate NES in TIRR have been given an identifier (NES1, NES2, NES3, and NES4) (**Figure 3B**).

**Figure 3:**
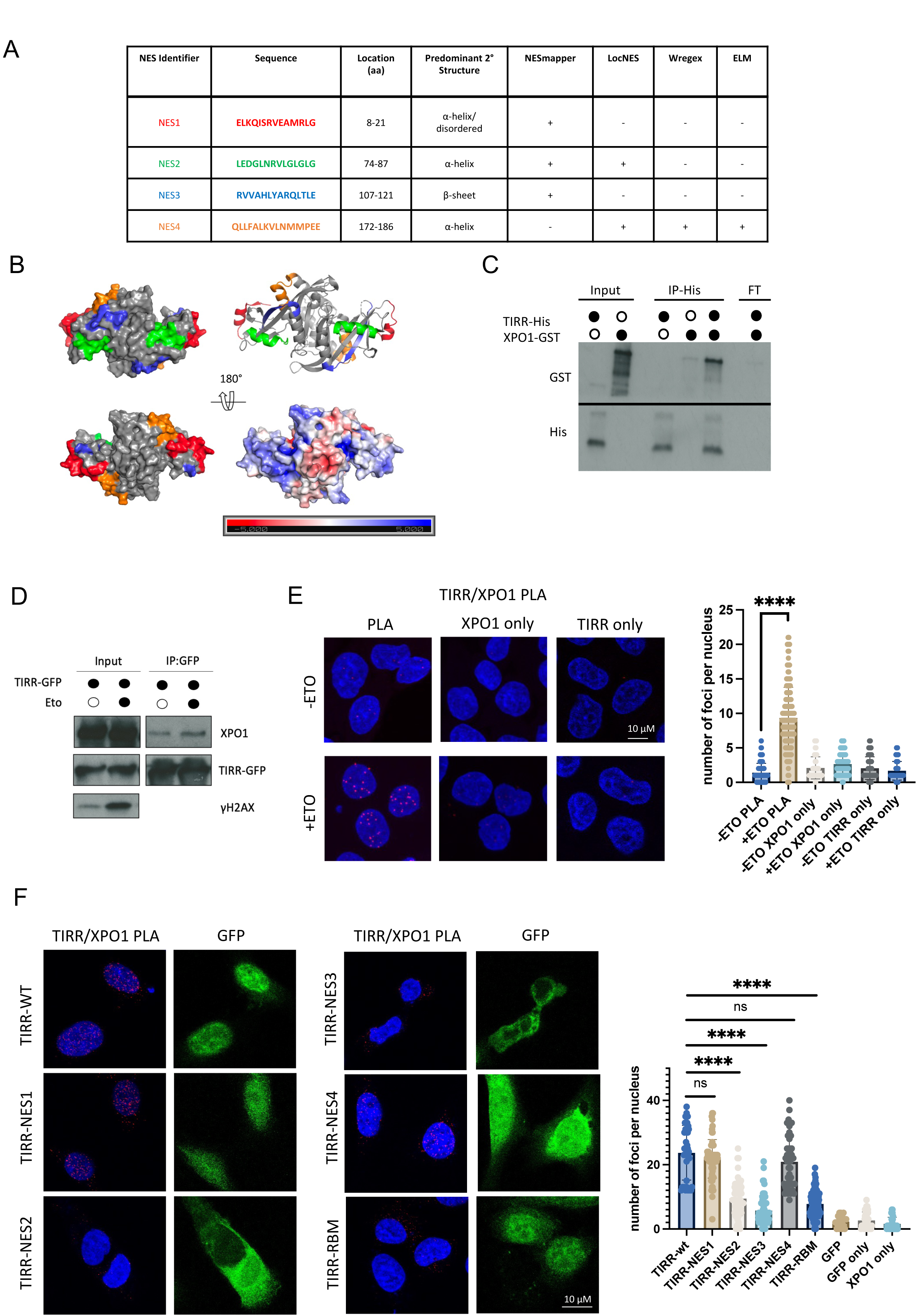
TIRR interacts with XPO1 upon DNA damage. A) Table showing predicted NES on TIRR. B) TIRR dimer (PDB: 6D0L) with the location of each predicted NES highlighted. The lower right structure shows the electrostatic landscape of TIRR, with the region displaying increased positive charge in blue indicating the RNA-binding region. C) *In vitro* interaction assay with purified TIRR-His and recombinant XPO1-GST. Immunoblot showing GST and His signals. D) Co-IP of TIRR-GFP and immunoblot for XPO1, GFP and gH2AX with and without Etoposide (ETO). E) Left: Confocal images showing PLA of TIRR and XPO1 in cells with or without ETO treatment. Single antibodies were used as negative control. Right: Quantification of PLA signals. Significance was determined using non-parametric Mann-Whitney test (****p ≤ 0.0001). F) Left: Confocal images showing PLA of TIRR-GFP wt, TIRR RNA binding mutant, and TIRR NES mutants in cells treated with Etoposide (ETO). Expression of GFP constructs is in green, PLA in red. DAPI indicates nuclei. Right: Quantification of PLA signals. Significance was determined using non-parametric Mann-Whitney test (****p ≤ 0.0001).

We next assessed whether TIRR can associate with XPO1. We incubated recombinant His-tagged TIRR (8) and GST-tagged XPO1, performed pulldown of TIRR and assessed for XPO1 binding. XPO1 was able to bind to TIRR in this assay, with minimal background binding of XPO1 detected on the beads in the absence of TIRR (**Figure 3C**). We then confirmed the interaction of TIRR and XPO1 through pulldown and western blot in both non-damage and damage conditions. By employing co-IP, we saw TIRR immunoprecipitated with XPO1 in both conditions, whilst no XPO1 was observed in GFP control pull down (**Figure 3D and Supplementary Figure S8C**). However, likely due to transient nature of this interaction, we were not able to observe a reproducible increase in this interaction in damage conditions. Therefore, we employed the quantifiable technique Proximity Ligation Assay (PLA), which is antibody based and allows for the detection of proximity between two proteins of interest in cells. PLA also allows the detection of weak and transient interactions. Using PLA with TIRR and XPO1 antibodies, we were also able to detect an interaction between TIRR and XPO1, which was significantly increased after DNA damage induced with either Etoposide (**Figure 3E**, single antibodies were used as a negative control), or IR (**Supplementary Figure S8D**). These data suggest that TIRR is an XPO1-interacting protein in the DDR.

NES sequences are relatively commonly found across the proteome, especially in hydrophobic cores of proteins, which in reality are not available for targeting and binding of XPO1 (43). True NES sequences are mainly found in accessible regions of proteins, in regions of alpha helical secondary structure, and are more likely to be within or adjacent to regions which are more flexible or locally disordered (43). Closer analysis of the identified candidate NES sequences revealed that candidates 1 and 3 contain regions of beta stranded structure (**Figure 3A and B**). However, it is not known whether structural changes may occur in TIRR upon RNA binding, and therefore we proceed with investigation of all four candidate NES sites.

We generated GFP-tagged mutants of all the NES sites by alanine scanning of hydrophobic residues at each of the NES sites, a method employed in validating NES sites in previous studies (44,45). We also generated a GFP-tagged TIRR RNA-binding mutant (26). Transfection of U2OS cells with plasmids corresponding to these mutants and subsequent PLA was performed in etoposide-treated cells. We found that mutation of the NES2 motif resulted in significantly decreased TIRR binding to XPO1 (**Figure 3F and Supplementary Figure S8E**). This motif is located distant to the RNA binding groove. Furthermore, TIRR binding to XPO1 was significantly reduced in cells expressing TIRR-NES3 mutant. NES3 motif is located internally in β-sheet structure near the RNA-binding groove. Mutations in NES1 and NES4 did not reduce TIRR binding to XPO1. Interestingly, TIRR RNA-binding mutant (TIRR-RBM) affected TIRR binding to XPO1. Expression of GFP alone served as a negative control. Also, single antibodies were used as a negative PLA control and did not result in any detectable foci (**Figure 3F and Supplementary Figure S8E**). These data suggest that TIRR binds to RNA and XPO1 via similar regions, or perhaps that RNA-binding is required for efficient XPO1 binding. All together, we conclude that TIRR binds the protein export factor XPO1 in part through direct interaction with one or more of its NES sequences.

### TIRR re-localizes from nucleus to cytoplasm upon DNA damage

Having confirmed TIRR associates with XPO1 upon DNA damage, we next examined the export of TIRR in damage and non-damage conditions, and whether this process depends on XPO1. In wildtype HeLa cells, we performed immunofluorescence of TIRR in control and ETO-induced damage conditions in the presence and absence of Leptomycin B (LMB), which inhibits XPO1 activity by preventing its nuclear reimport (46). In the absence of LMB, we observed a significant increase in localization of TIRR to the cytoplasm upon DNA damage. Upon the addition of LMB, TIRR was retained in the nucleus, suggesting TIRR export is mediated by XPO1 (**Figure 4A**). To validate our data further, we performed the same experiment in cells treated with IR to induce DNA damage and observed damage-dependent, LMB-sensitive, TIRR re-localisation into the cytoplasm (**Supplementary Figure S9A).** To eliminate the possibility that this phenotype is cell-type specific, we also performed this IF experiment in U2OS cells treated with ETO and detected damage-induced TIRR localisation to the cytoplasm, which was inhibited by LMB treatment (**Supplementary Figure S9B).** To test the nuclear to cytoplasmic shift in real time, we employed a previously characterised engineered photoactivatable (PA) variant of GFP, which increases in fluorescence 100 times after irradiation with 413 nm light and allows for protein tracking *in vivo* (47). We generated a photoactivatable GFP-tagged TIRR construct (PAGFP-TIRR) and transfected U2OS cells with both PAGFP-TIRR plasmid and a mCherry-LaminB1-10 plasmid, chosen to delineate the borders of nuclei (**Figure 4B, illustration and Supplementary Figure S9C**)(25). To induce DNA damage, IR was chosen as it induces damage on a shorter, more concentrated time scale than ETO, allowing us to observe TIRR re-localization very quickly after damage induction. A combined IR damage induction and LMB treatment condition was also included. PAGFP was activated with 413 nm light at a single point within the nucleus followed by time lapse imaging for 120 seconds. In no damage conditions, GFP signal did not increase significantly in the cytoplasm over time (**Figure 4B and Supplementary movie 1**). However, in damage conditions, GFP signal in the cytoplasm increased significantly (**Figure 4B and Supplementary movie 2**). In the LMB treated damage condition, no detectable increase in cytoplasmic GFP signal was observed (**Figure 4B and Supplementary movie 3**), further confirming the importance of XPO1 in facilitating export. These data suggest that DNA damage stimulates LMB-sensitive TIRR export from nucleus into cytoplasm.

**Figure 4:**
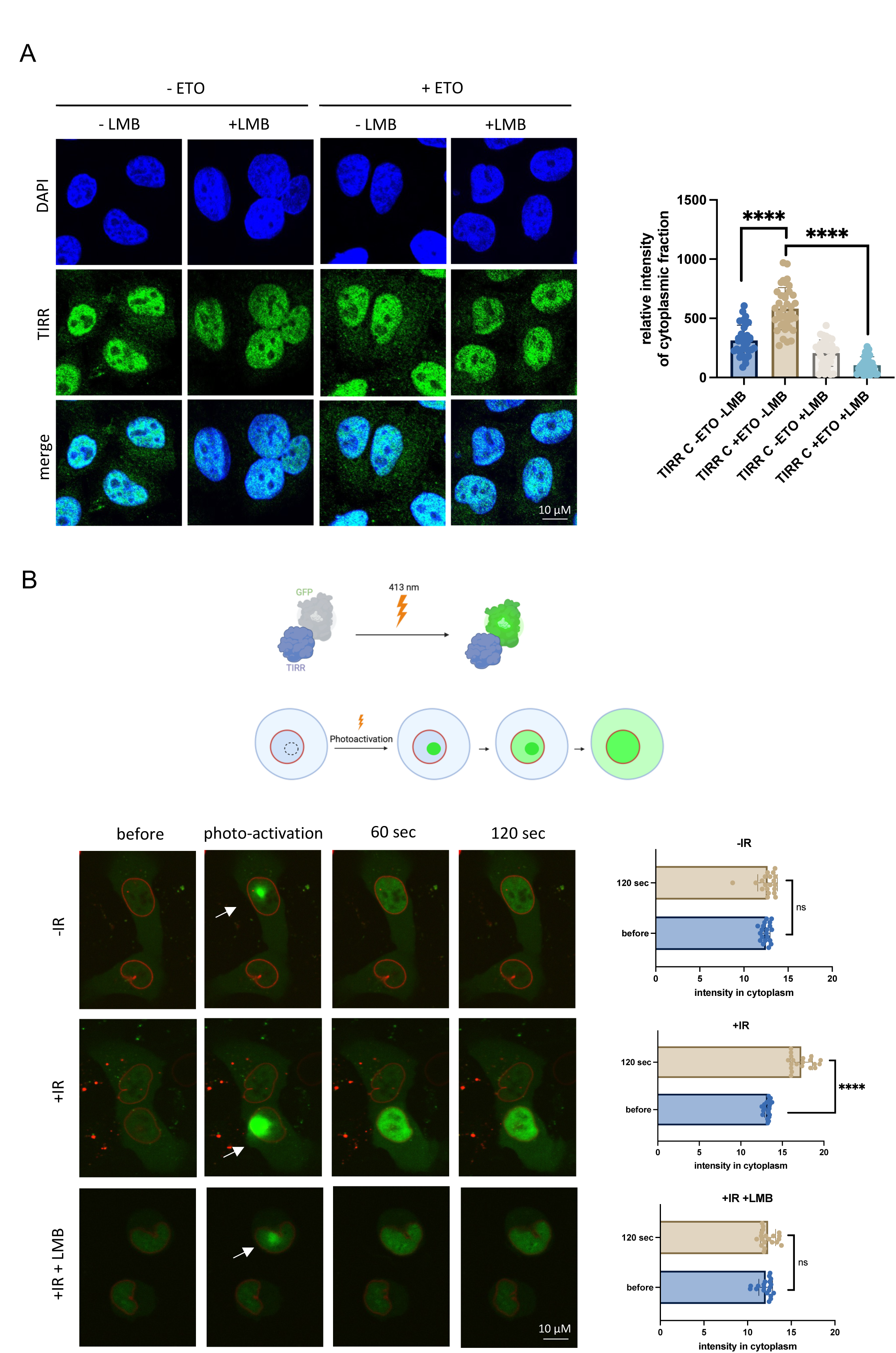
TIRR re-localizes from nucleus to cytoplasm upon DNA damage. A) Left: Representative confocal images showing immunofluorescence of TIRR in HeLa cells with or without Etoposide (ETO) treatment and with or without LMB treatment. DAPI was used to stain nuclei. Right: Quantification of TIRR signal in cytoplasm. n>50 cells, significance was determined using t- test, (****p ≤ 0.0001). B) Top: diagram showing photoactivation strategy. Bottom: representative confocal images showing live cells expressing PAGFP-TIRR (in green), mCherry-LaminB1-10 was used as a marker for the nuclear envelope. Photoactivation was induced using 413 nm laser and the location of activation is indicated with a white arrow. Photoactivation was followed by time lapse microscopy, with time points at 60 and 120 seconds shown. Quantification of cytoplasmic GFP signal was performed using FIJI software. Significance was determined using t- test, (****p ≤ 0.0001). See also Supplementary movies 1-3. Illustration made with Biorender.com.

### TIRR associates with and regulates the formation of P-bodies in response to DNA damage

Analysis of RIP-seq data revealed that TIRR binds preferentially to RNAs with a lower GC content (**Figure 1F**), which have been shown to be enriched in P-bodies (18). Interestingly, PBs are considered both sites of RNA decay and storage, influencing the decay and stability of specific RNAs (17,18). RNAs targeted to PBs appear to be related to regulatory processes such as RNAPII transcription allowing PBs to dynamically influence the cellular response to specific stimuli (24). Therefore, we wondered whether TIRR and its bound RNA could be associated with PBs. To test this hypothesis, we performed co-immunofluorescence (co-IF) of TIRR and LSM14A (LSM14A is an established marker of P-bodies and is essential for their formation (24,48,49)) using antibodies against endogenous proteins and observed distinct TIRR foci colocalizing with LSM14A foci. This co-localisation was significantly increased upon DNA damage, as assessed by Pearson’s co-localisation coefficient (**Figure 5A).** Next, we overexpressed TIRR in HeLa cells and performed co-IF using antibodies against GFP and LSM14A in cells subjected to IR. Distinct foci, containing overlapping green (TIRR-GFP) and red (LSM14A) signal were detected in damaged cells (**Supplementary Figure S10A).** We also employed Co-IP of TIRR and LSM14A in TIRR-GFP cells and revealed that LSM14A was pulled down by TIRR (**Figure 5B**). To test the interaction between TIRR and LSM14A in live cells, we transfected U2OS cells with PAGFP-TIRR and LSM14A-mScarlet and observed interaction between these two proteins in the form of overlapping foci that co-localised continuously within the tested time frame (**Figure 5C and Supplementary movie 4**).

**Figure 5:**
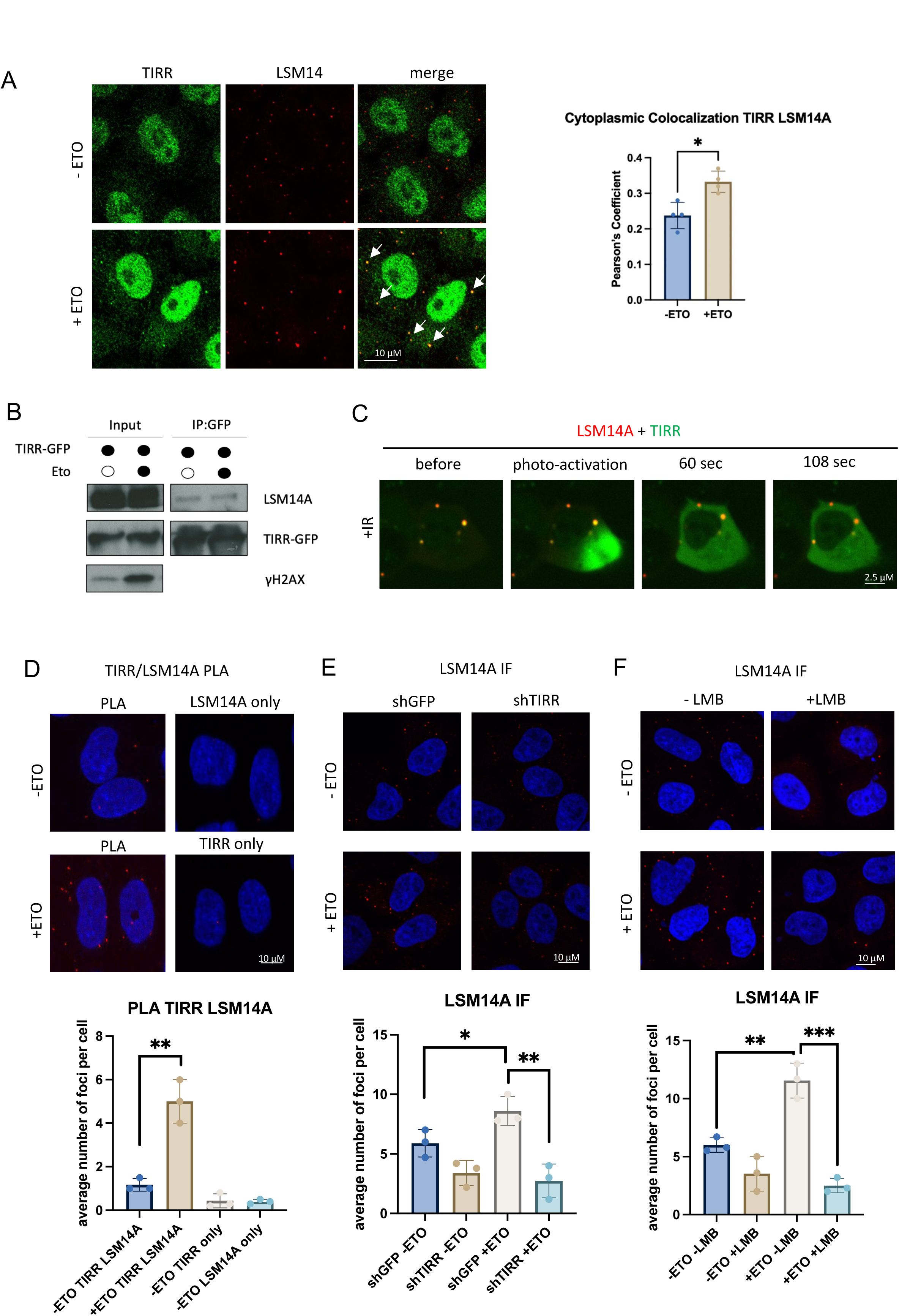
TIRR associates with and regulates P-bodies formation in response to DNA damage. A) Left: Representative confocal images showing immunofluorescence of TIRR in cells with or without Etoposide (ETO) treatment. TIRR expression was visualised in green channel (GFP), P-bodies were labelled using anti-LSM14A antibody and visualised in red channel. White arrows point to P-bodies. Right: Quantification of TIRR and LSM14A co-localisation signal in cytoplasm using Pearson’s coefficient, n>50 cells, significance was determined using t- test, (*p ≤ 0.05). B) Co-IP of TIRR-GFP and immunoblot for LSM14A, GFP and gH2AX with and without Etoposide (ETO). C) Representative confocal images showing live cells expressing PA-TIRR-GFP (in green) and LSM14A-mScarlet (in red) as a marker for P-bodies. Photoactivation was induced using a 413 nm laser, and the location of activation is indicated with a white arrow. Photoactivation was followed by time lapse microscopy, time points at 60 and 108 seconds are shown. See also Supplementary movie 4. D) Top: proximity ligation assay using anti-TIRR and anti-LSM14A antibodies in no damage (-ETO) or damage (+ETO) conditions in HeLa cells. DAPI was used to label nuclei. PLA signals were visualised in red channel. Single antibodies were used as negative controls. Bottom: quantification of top. Error bar = mean ±SD, significance was determined using non- parametric Mann Whitney test (**p ≤ 0.01). E) Top: Immunofluorescence analysis showing LSM14A staining in no damage (-ETO), and damage (+ETO) conditions in cells expressing either shGFP (control) or shRNA targeting TIRR (shTIRR). DAPI was used to stain nuclei. Bottom: Quantification of TIRR signal in cytoplasm, n>50 cells, significance was determined using t- test, (**p ≤ 0.01, *p ≤ 0.05). F) Top: Immunofluorescence analysis showing LSM14A staining in no damage (-ETO), and damage (+ETO) and with or without LMB treatment. DAPI was used to stain nuclei. Bottom: Quantification of TIRR signal in cytoplasm, n>50 cells, significance was determined using t- test, (***p ≤ 0.001, **p ≤ 0.01).

Finally, we employed PLA using antibodies against TIRR and LSM14A in HeLa and U2OS cells to assess whether their visual proximity was reflected in molecular proximity. Indeed, upon ETO induced DNA damage in both cell lines, PLA foci number increased significantly, suggesting proximity (<∼40nm) of TIRR and LSM14A after DNA damage (**Figure 5D and Supplementary Figure S10B**). TIRR and LSM14A antibody only in non-damage conditions were used as negative controls. Similarly, we observed damage dependent PLA foci formation between TIRR and LSM14A in HeLa cells subjected to IR (**Supplementary Figure S10C).** Collectively, we have shown that TIRR associates with P bodies, and this interaction is enhanced by DNA damage.

To further understand the significance of this interaction, we performed immunofluorescence of LSM14A foci in shTIRR and shGFP HeLa cells with induction of DNA damage by ETO or IR. Upon DNA damage in control conditions, an increase in P body number was observed when compared to non-damage. This is in agreement with previous studies of P body formation, where it was found that various stresses and DNA damage conditions resulted in increased P body formation (50–53). Interestingly, TIRR depletion resulted in decreased P body formation (**Figure 5E and Supplementary Figure S10D**). We show that LSM14A protein levels are not affected in TIRR KD cells (**Supplementary Figure S10E),** suggesting that TIRR is modulating P body formation via a different mechanism. We also show that the overexpression of TIRR- GFP could elicit an increase in P body numbers (**Supplementary Figure S10F-G**). These data suggest that TIRR can colocalize with P bodies, in particular after damage, and can modulate P body formation.

We showed that TIRR can interact with XPO1. Therefore, we assessed whether TIRR modulation of P body numbers is dependent on XPO1 activity. We performed immunofluorescence of LSM14A upon damage and inhibition of XPO1 with LMB. Indeed, P body number was significantly reduced in DNA damage induced by ETO upon LMB treatment in HeLa or U2OS cells **(Figure 5F and Supplementary Figure S11A)**. Similarly, this was also observed in HeLa cells subjected to IR (**Supplementary Figure S11B).** Therefore, we conclude that TIRR can regulate the formation of P bodies, and this function is dependent on XPO1.

### TIRR bound mRNAs are associated with P bodies

We detected TIRR being associated with P bodies (**Figure 5**) and therefore we wished to investigate whether TIRR bound mRNA is also localised to PBs. First, we performed RNA Fluorescence in situ hybridisation in the presence or absence of ETO with probes against *ZNF600* and *PELI2*, two mRNAs found to bind to TIRR specifically upon damage in our RIP-Seq, as well as probes against *SPEN*, an mRNA known to localize to P bodies but not found in our RIP-Seq, and *Tubulin*, an mRNA also not found in our RIP-Seq (24). This was followed by immunofluorescence co-staining with antibody for LSM14A in HeLa cells expressing either shTIRR or shGFP. Close analysis of confocal images revealed assembly of *ZNF600* or *PELI2* FISH probes in cytoplasmic foci (green) that overlapped with LSM14A (red) IF signals. This co-localisation, assessed by Pearson co-localisation coefficient, was abolished in TIRR KD cells, most likely due to impaired P body formation and mRNA nuclear retention (**Figure 6A and B**). Additionally, *SPEN* and *Tubulin* colocalization with P bodies in damage conditions remained apparent and was not affected by depletion of TIRR (**Figure 6C and D**). FISH- LSM14A foci of *ZNF600* or *PELI2* were also observed in cells subjected to IR (**Supplementary Figure S11C and D**).

**Figure 6:**
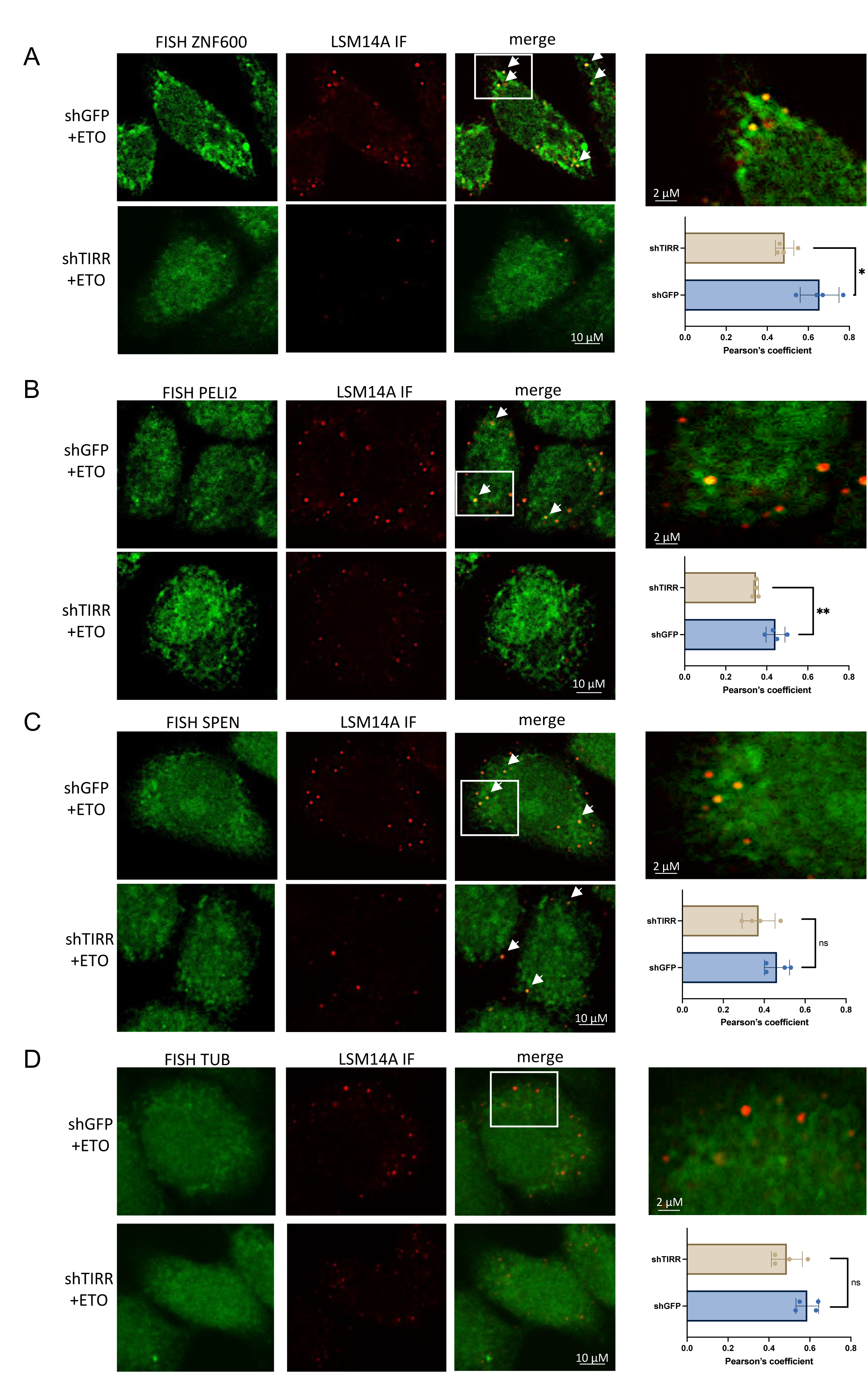
TIRR depletion leads to accumulation of mRNA in nucleus. A) Representative confocal images showing co-localisation of RNA FISH signals (in green) for *ZNF600* with LSM14A (in red) in shGFP or shTIRR cells after ETO treatment. n>50 cells. White rectangles mark zoomed in images on the right. White arrows point to P-bodies with FISH signal (in yellow). Quantification of FISH and LSM14A co-localisation signal using Pearson’s coefficient, n>50 cells, significance was determined using t- test, (*p ≤ 0.05, **p ≤ 0.01). B) As in A) for *PELI2* mRNA. C) As in A) for *SPEN* mRNA D) As in A) for *Tubulin* mRNA

These data show that TIRR bound mRNAs are localised in PBs upon DNA damage in a TIRR- dependent manner.

### The role of TIRR in mRNA export contributes to efficient DNA repair

TIRR has been identified as a negative regulator of 53BP1. The complex of 53BP1/TIRR is constitutive in non-damage conditions, however after DSB induction, the complex dissociates, allowing 53BP1 to recognize and bind to chromatin to promote NHEJ (8). Several studies showed that depletion of TIRR sensitises cells to induced DNA damage (5,54,55). This effect was solely attributed to TIRR’s role in 53BP1 regulation, and possible implications of its role in RNA binding has largely been omitted. To investigate whether TIRR’s role in mRNA export has any impact on DNA repair, we assessed the interaction of 53BP1 and the NES mutants and TIRR-RBM by performing co-IP of TIRR-GFP and 53BP1 in U2OS cells transfected with corresponding plasmids in ETO-induced damage conditions. We detected TIRR-WT binding to 53BP1 even in damage conditions, as observed previously (8) (**Figure 7A**). Intriguingly, while the interaction appears somewhat reduced, TIRR-RBM and TIRR-NES1 and TIRR- NES2 were found to retain 53BP1 binding ability (**Figure 7A**). Therefore, we were interested in whether there may be 53BP1-binding independent effects of these mutants on DNA damage clearance efficiency. In U2OS cells, we performed endogenous TIRR knockdown and subsequent overexpression of TIRR-WT, TIRR-K10E (a well-characterized 53BP1-binding mutant (4,54,56), TIRR-RBM and TIRR-NES2, as well as an GFP only plasmid control (**Supplementary** Figure 12A). DNA damage was induced by 10 Gy IR and immunofluorescence of γH2AX was performed on samples collected at 15 minutes, 6 hours, and 24 hours after damage induction. After 6 hours, TIRR-K10E, TIRR-RBM and TIRR-NES2 all displayed significantly increased γH2AX intensity relative to TIRR-WT rescue, suggesting reduced γH2AX clearance efficiency (**Figure 7B**). At 24 hours post-damage induction, again all three mutants had significantly greater γH2AX intensity relative to WT rescue, with TIRR- RMB having the most significant difference (**Figure 7C**). Therefore, TIRR may have other contributions to the efficiency of the DDR beyond 53BP1 binding, likely through its RNA-binding and export abilities.

**Figure 7:**
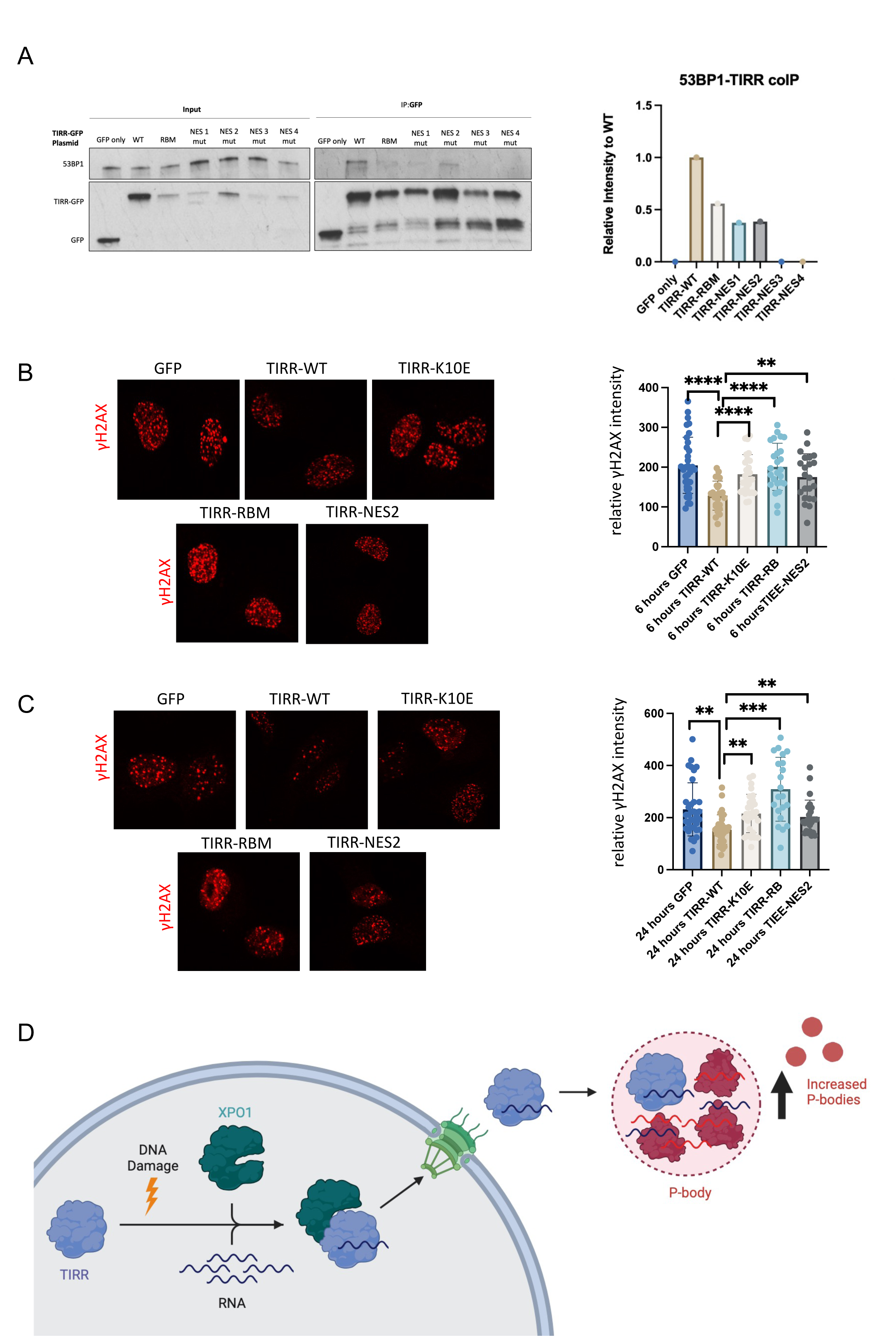
TIRR facilitates mRNA export and is associated with P-bodies upon DNA damage. A) Left: Co-IP of 53BP1 with TIRR mutants in ETO-induced damage conditions. U2OS cells were transfected with either GFP, TIRR-WT, TIRR-RMB TIRR-NES1, TIRR-NES2, TIRR- NES3, or TIRR-NES4. Right: Normalization of samples to TIRR-GFP pulldown. B) Immunofluorescence showing γH2AX signal in cells transfected with siTIRR for endogenous TIRR KD, followed by transfection with GFP, TIRR-WT, TIRR-K10E, TIRR- RBM and TIRR-NES2 mutant, at 6 hours post IR treatment. Significance was determined using t-test, (**p ≤ 0.01, ****p ≤ 0.0001). C) Immunofluorescence showing γH2AX signal in cells transfected with siTIRR for endogenous TIRR KD, followed by with GFP, TIRR-wt, TIRR-K10E, TIRR-RBM and TIRR- NES2 mutant, at 24 hours post IR treatment. Significance was determined using t- test, (**p ≤ 0.01, ****p ≤ 0.0001). D) Model: TIRR binds specific mRNAs and mediates their nuclear export through XPO1. In cytoplasm, TIRR sequesters mRNA into P-bodies, in order to stabilise them during damage conditions. Image was created with BioRender.

## Discussion

The RNA binding function of TIRR, particularly in the DDR, remains largely unexplored. In this study, we demonstrate that TIRR exhibits binding affinity towards a specific subset of mRNAs encoding genes involved in RNAPII regulatory processes, including transcription factors (**Figure 1D and E**). Notably, these mRNAs exhibit a significantly lower GC content compared to the rest of the genome (**Figure 1F**). Much like other NUDIX proteins, such as DCP2, TIRR can localise to P bodies, particularly following DNA damage, and has the capability to regulate PB numbers. Considering the observed impact of TIRR on PBs formation, we propose that TIRR selectively binds to a subset of mRNAs to facilitate their sequestration into PBs. Consistent with this hypothesis, an analysis of RNAs enriched in PBs revealed their involvement in regulatory pathways that necessitate dynamic regulation, such as RNA-processing, cell division, and RNAPII transcription(24). Our data demonstrate that TIRR localises subset of mRNAs into PBs, particularly after DNA damage (**Figure 6**).

In our work we identified a previously unknown interaction of TIRR with XPO1 and characterise candidate NES sequences within TIRR (**Figure 2A-B**). Over 1000 proteins in human cells are suggested to be regulated by XPO1-dependent export (42), including proteins involved in translation and mRNA degradation (42). TIRR is both a nuclear and cytoplasmic protein (5,6), with functions in both compartments. We show that TIRR interacts with XPO1 after DNA damage (**Figure 3**) and identify two specific binding motifs, NES2 and NES3, required for direct binding to XPO1. Interestingly, mutation of TIRR’s RNA-binding domain affects its interaction with XPO1, suggesting that association of TIRR with RNA may enhance XPO1 binding. We also show a portion of TIRR is exported from the nucleus after damage via an XPO1-mediated pathway (**Figure 4**).

Analysis of RNA-seq did not reveal large changes in total RNA levels in response to TIRR knockdown and ETO at the steady state level, nor did we detect significant changes in protein product levels of several transcripts bound to TIRR upon damage. Microscopically visible P bodies have not conclusively been shown to be required for activities such as mRNA decay and miRNA-mediated translational silencing, nor does targeting of an mRNA to P bodies necessitate degradation (21,48,57). Yeast cells deficient in *lsm4* and *edc3* genes do not form P bodies, and yet still show mRNA decay activity (58). It has been posited that P bodies may only form due to the accumulation of translationally repressed mRNA complexes, but do not themselves instigate decay or silencing (48). TIRR has previously been found to inhibit translation in non-damage conditions, thus it may perform a similar role in damage conditions, contributing to the pool of translationally repressed mRNAs to scaffold P body formation (6). However, other roles of P bodies have been suggested, including a storage function for inactive mRNA decay factors and translationally repressed transcripts and/or maintenance of the stability of localized mRNA through mechanisms such as polyA protection (58,59).

We show that TIRR binds to mRNAs involved in dynamic regulatory processes, such as transcription, particularly after DNA damage (**Figure 1**). Part of cellular response to DNA damage is decreased transcription (60). Therefore, it is possible that TIRR selectively sequesters mRNA transcription factor transcripts to RNA granules for their storage, indirectly contributing to DSB induced transcriptional repression. As we have not directly assessed decay intermediates of TIRR-bound mRNA in DNA damage, we cannot exclude that some modulation of mRNA metabolism is occurring and is regulated by TIRR. Beyond P body mediated processes, in our model, these mRNA originate in the nucleus, where most decay of mature mRNA transcripts and translation does not occur (61,62). It is therefore within expectations that RNA and protein levels would largely remain unchanged during nuclear retention after TIRR knockdown.

Rather than TIRR bringing mRNA to the cytoplasm, it may be more important that TIRR is removing mRNA from the nucleus. Export of specific mRNA has been implicated in DDR previously. Studies in fertilized *Xenopus laevis* eggs showed that depletion of Gle1, a cofactor required for efficient Ddx19 mediated mRNA export, or Nxf1, a factor associated with the export of mRNA, induced DNA damage as detected by yH2AX foci (63,64). DNA damage induced by a DNA Topoisomerase I inhibitor [camptothecin (CPT)] was at least partially dependant on mRNA export for resolution. TIRR depletion does indeed increase yH2AX foci in non-damage conditions, however this has only been studied in the context of 53BP1 binding (5). It is therefore possible that TIRR may have a role in DNA repair through depletion of specific transcripts from the nucleus. Indeed, we show that TIRR depletion and rescue with either TIRR mutated at its RNA-binding domain or at the NES2 NES site, both of which retain 53BP1 binding, are insufficient to efficiently clear DNA damage as detected by γH2AX foci (**Figure 7A-B**). We also show that this effect is comparable or even more deleterious than rescue with the TIRR-K10E 53BP1-binding mutant (**Figure 7A-B**).

Overall, here we identified TIRR as a regulator of mRNA localization in response to DNA damage. TIRR selectively binds to a subset of mRNA encoding for proteins involved in transcription regulation, relocates to the cytoplasm through XPO1, and sequesters RNA into P bodies (**Figure 7D**). We also suggest that this function is important for efficient DNA damage repair. These findings extend our understanding of RNA regulation upon DNA damage.

## Data availability

All data are available in the paper and the Supplementary Data. The raw and processed RIP-seq and RNA-seq datasets described in this manuscript are deposited under GEO accession number GSE227686 and accessible with token: avchqsiutdyhjab (https://www.ncbi.nlm.nih.gov/geo/query/acc.cgi?acc=GSE227686).

## Acknowledgements

We would like to thank all the members of the Gullerova lab for their help and advice throughout this study. We would also like to thank Franca Esposito, Rosario Avolio, and Danilo Swann Matassa for their helpful discussions on TIRR and for providing the TIRR-GFP, GFP, shGFP and shTIRR cell lines. We are grateful to Alan Wainman for his help with microscopy.

## Author contributions

M.S.G. performed most of the experiments. I.C. performed all of the bioinformatics analysis. A.A. designed and performed the FISH experiments. R.F.K. performed some of the experiments and wrote parts of the manuscript. M.S.G. wrote the draft of the manuscript. M.G. designed and supervised the project and edited the manuscript, and assisted with microscopy. All the authors reviewed and approved the final version of the manuscript.

## Materials and correspondence

All correspondence should be addressed to M.G.

---

**Supplementary Figure 1.**
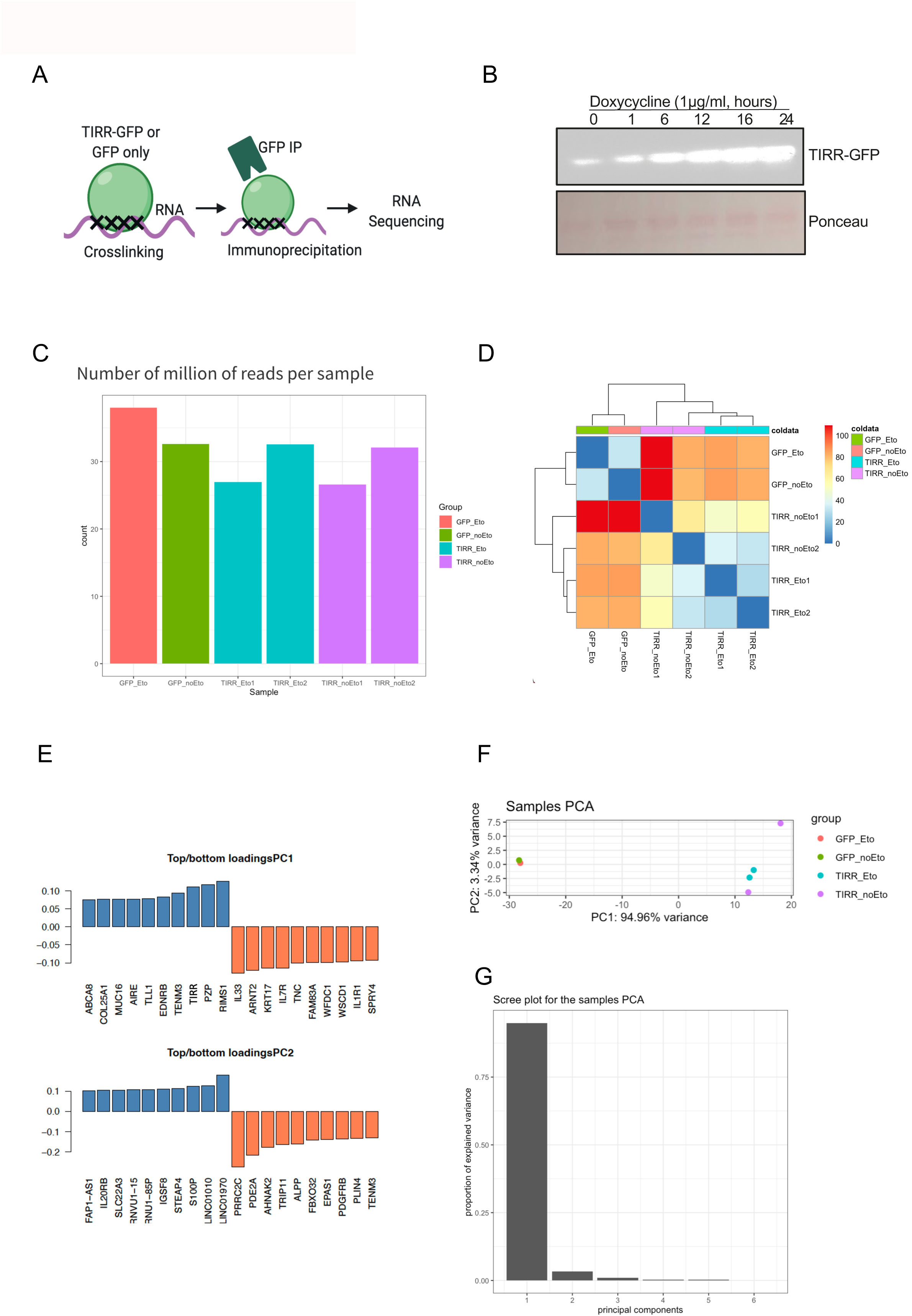

**Supplementary Figure 2.**
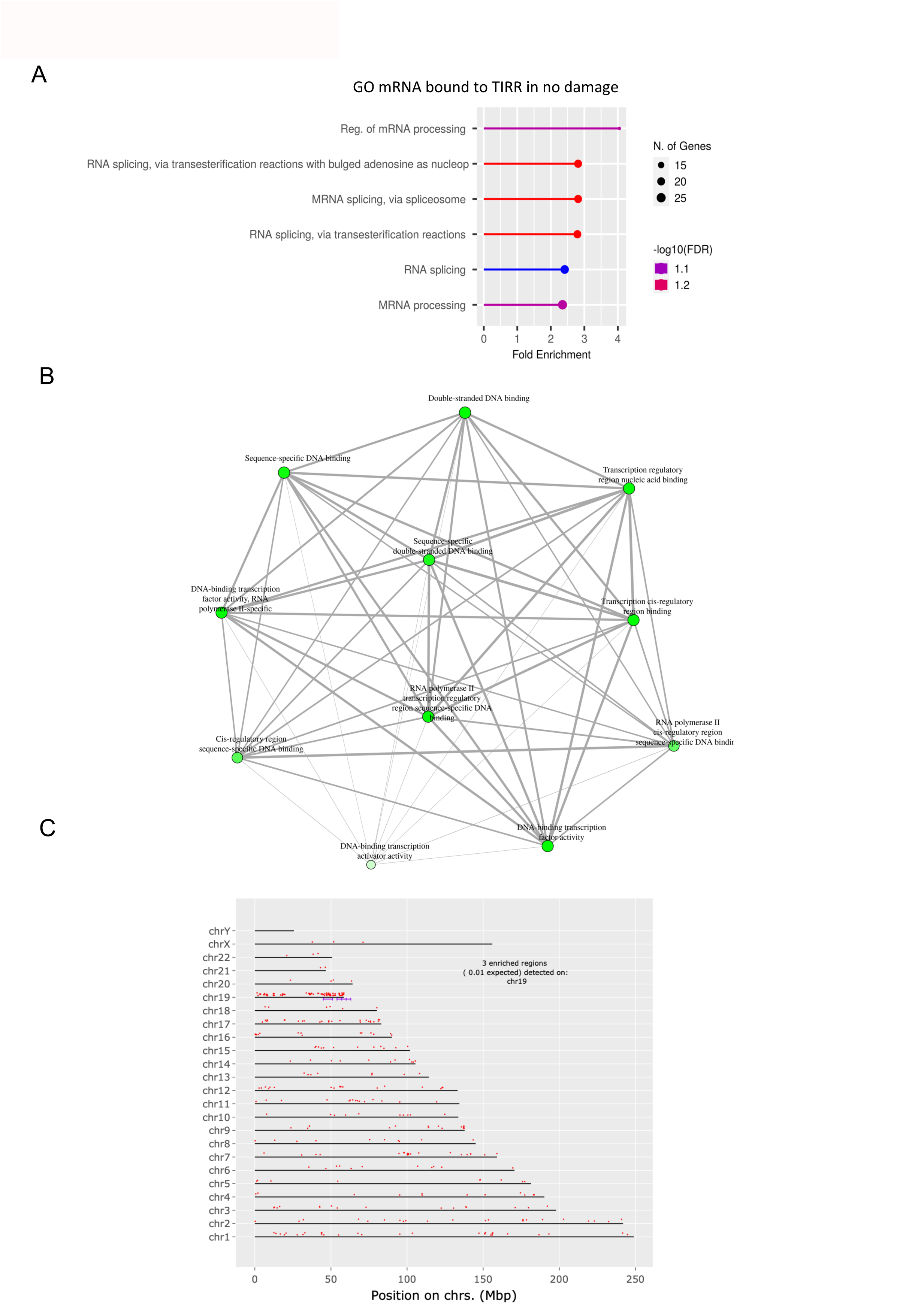

**Supplementary Figure 3.**
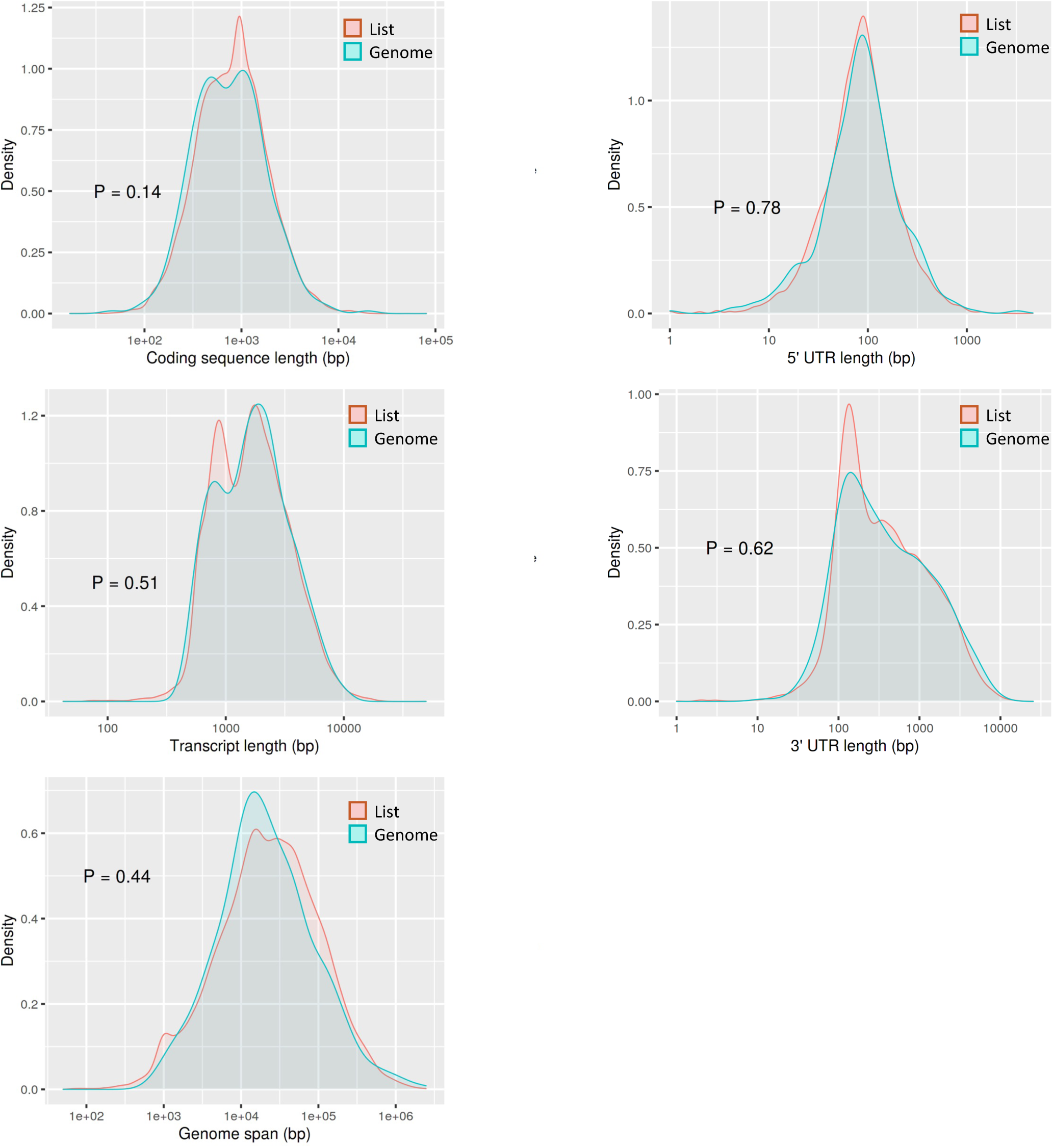

**Supplementary Figure 4.**
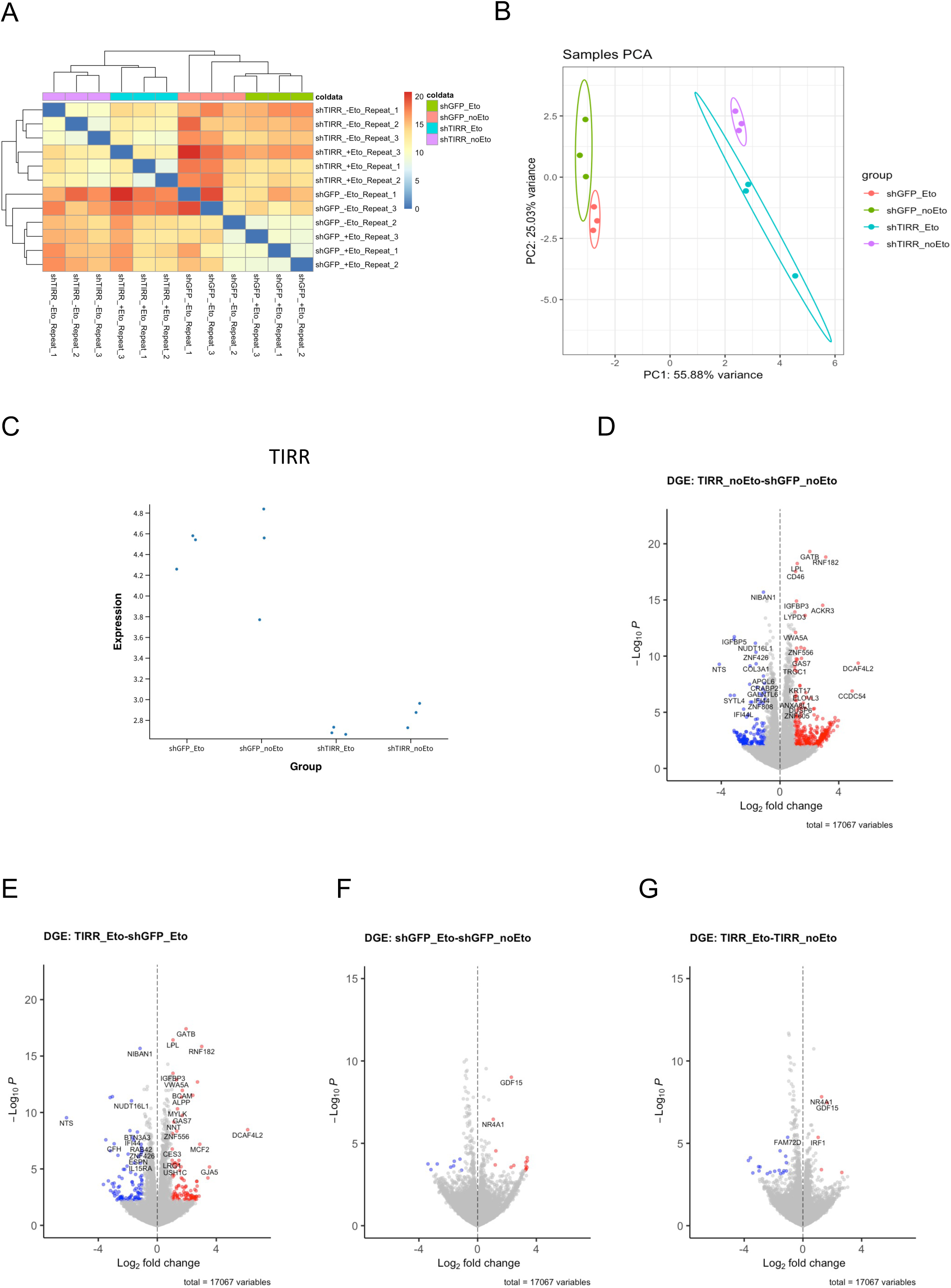

**Supplementary Figure 5.**
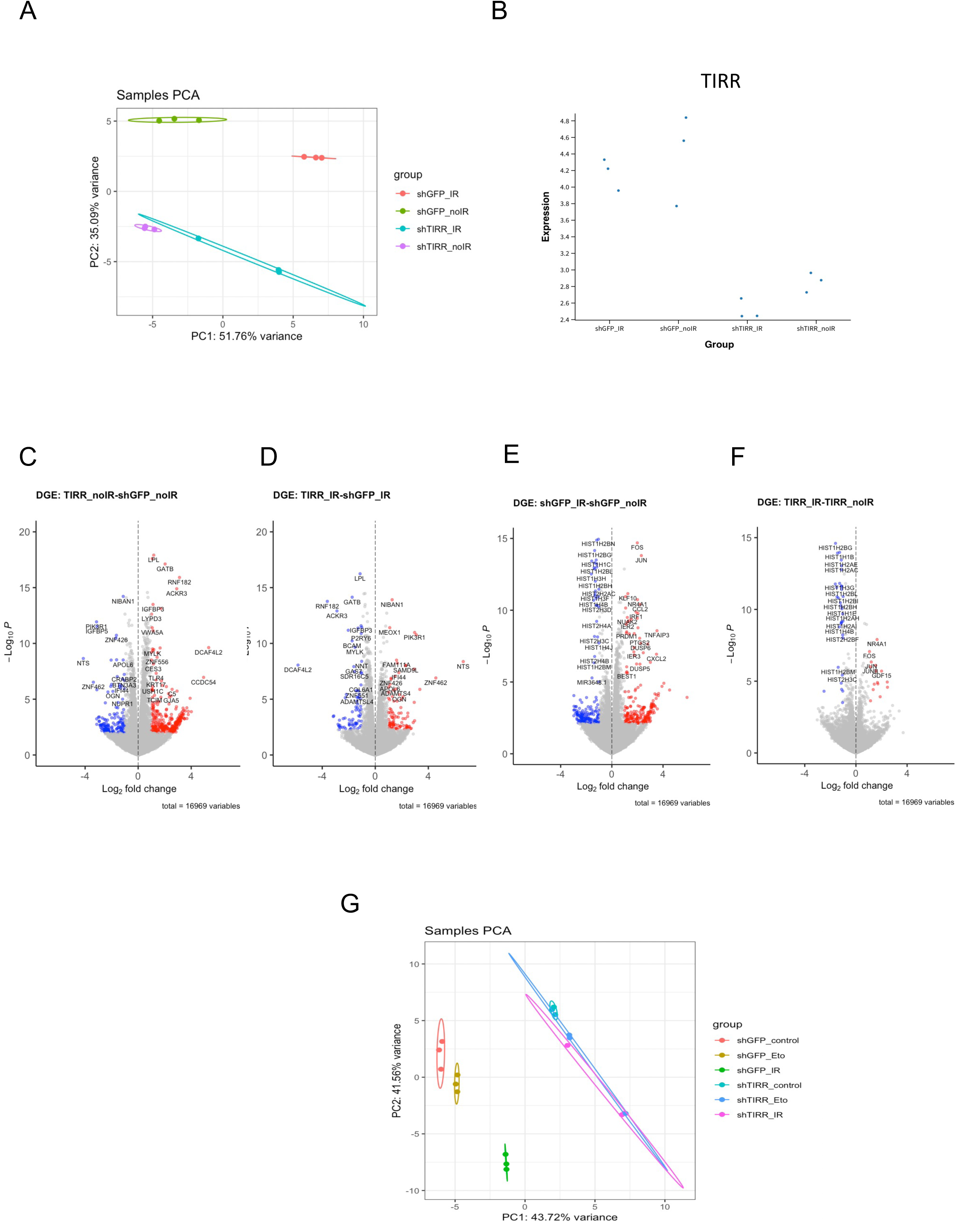

**Supplementary Figure 6.**
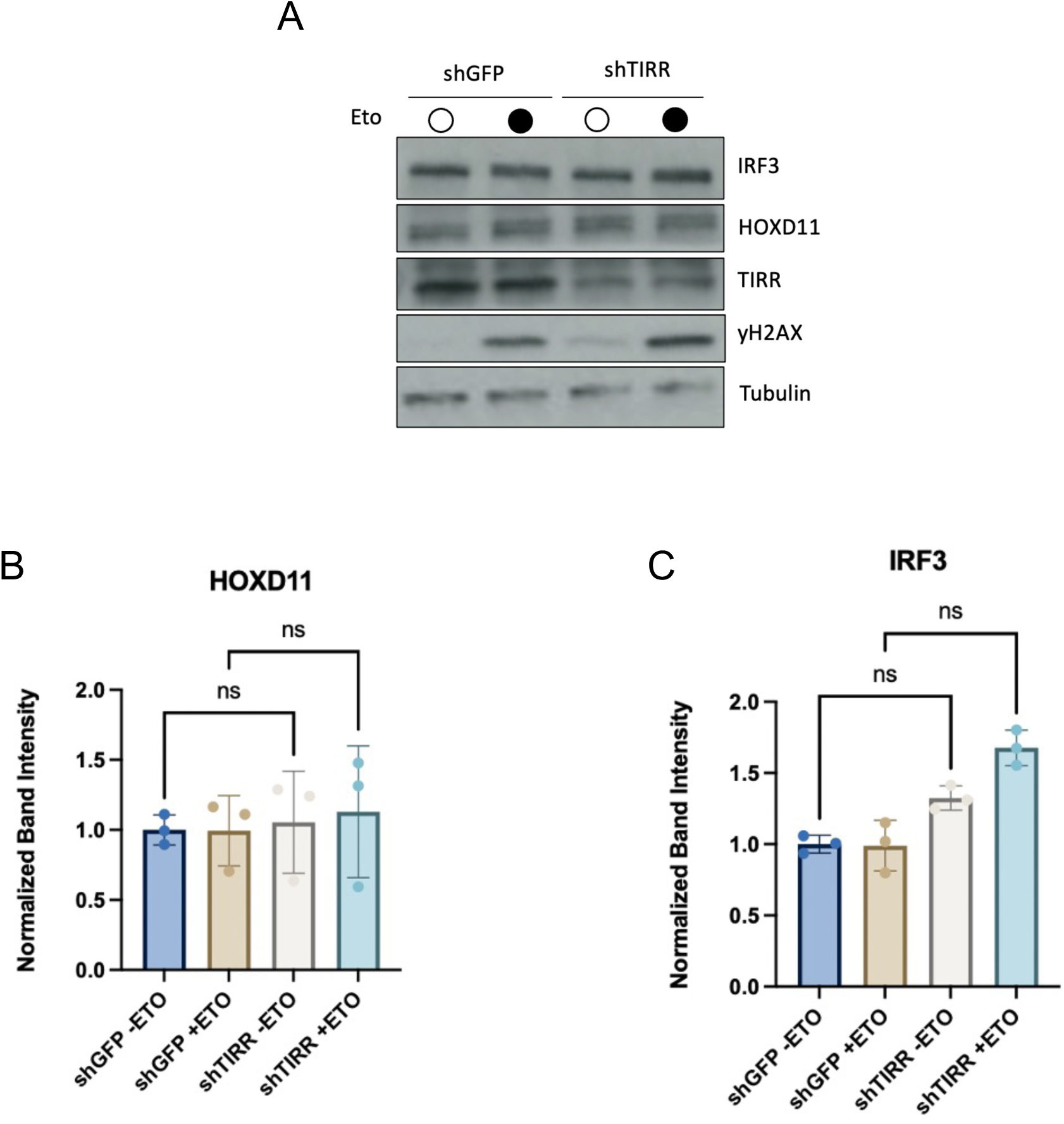

**Supplementary Figure 7.**
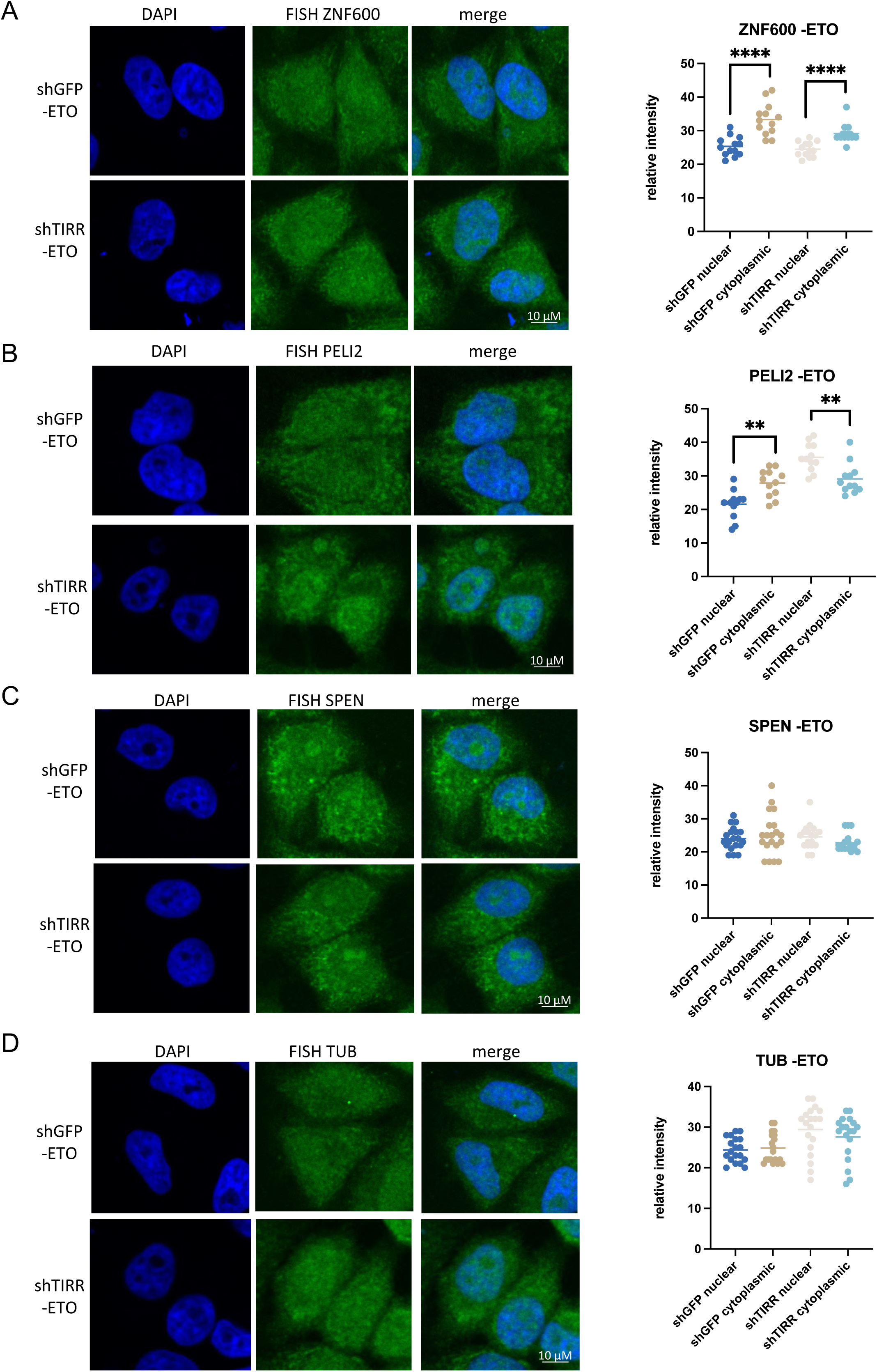

**Supplementary Figure 8.**
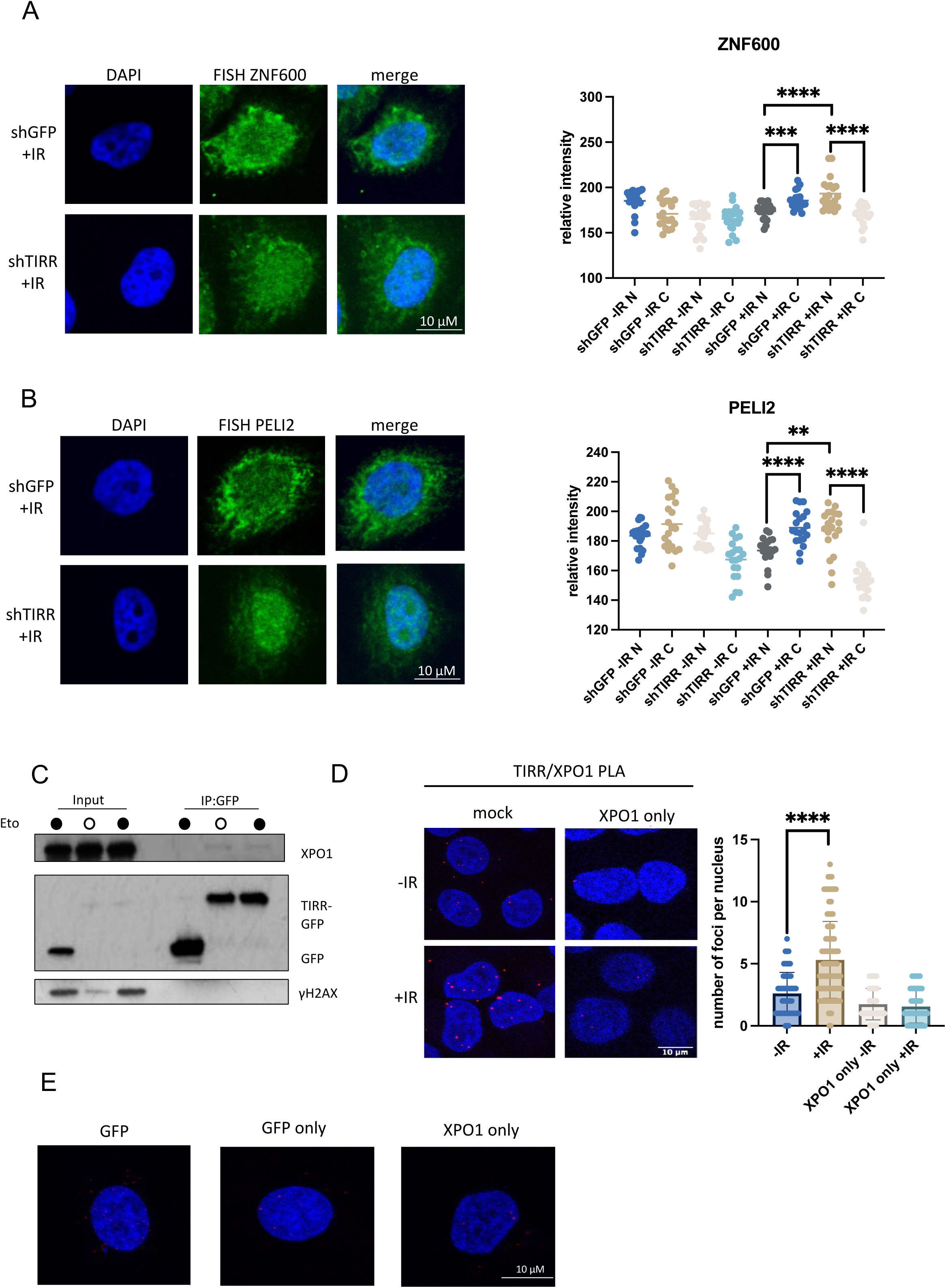

**Supplementary Figure 9.**
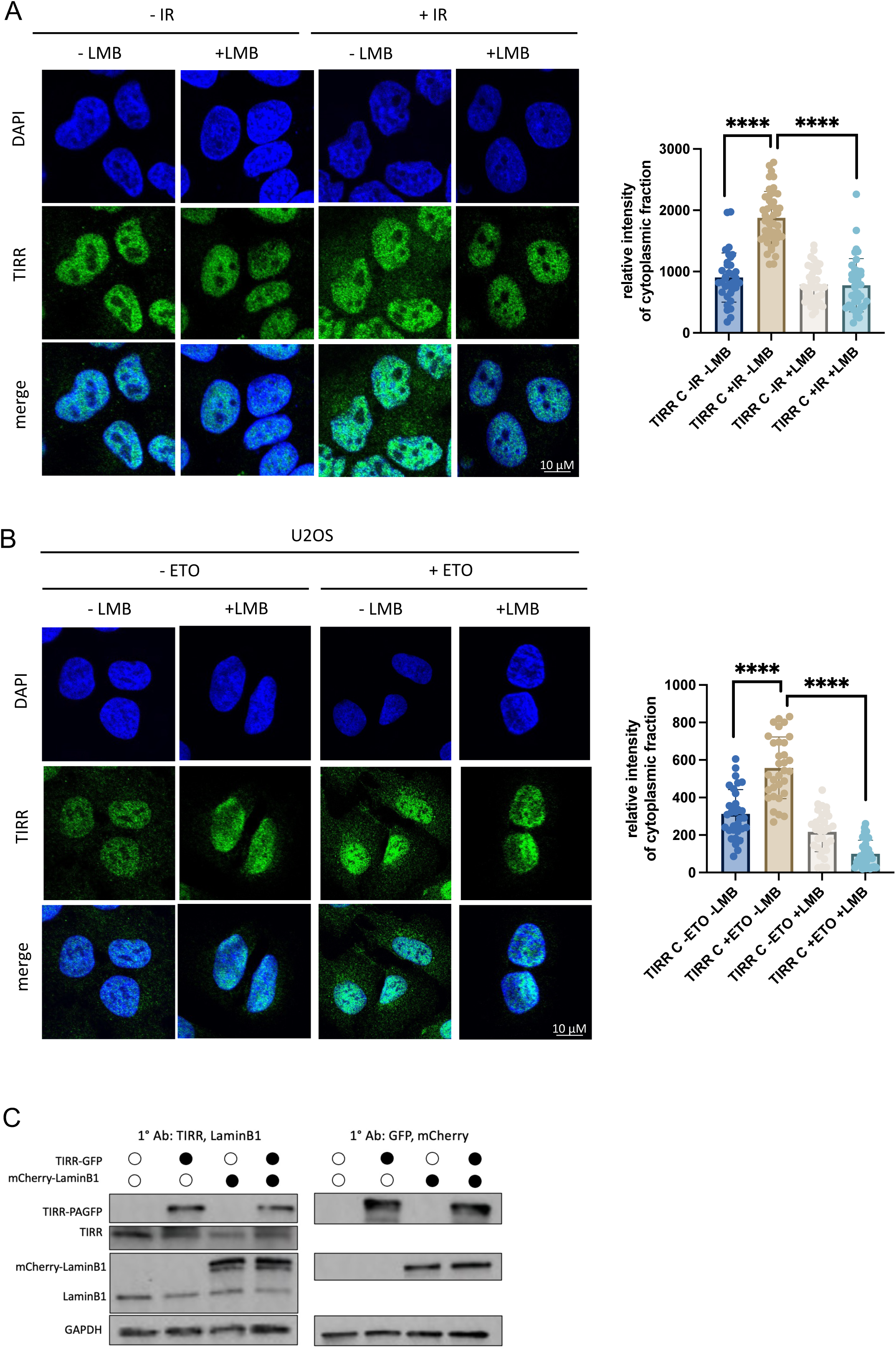

**Supplementary Figure 10.**
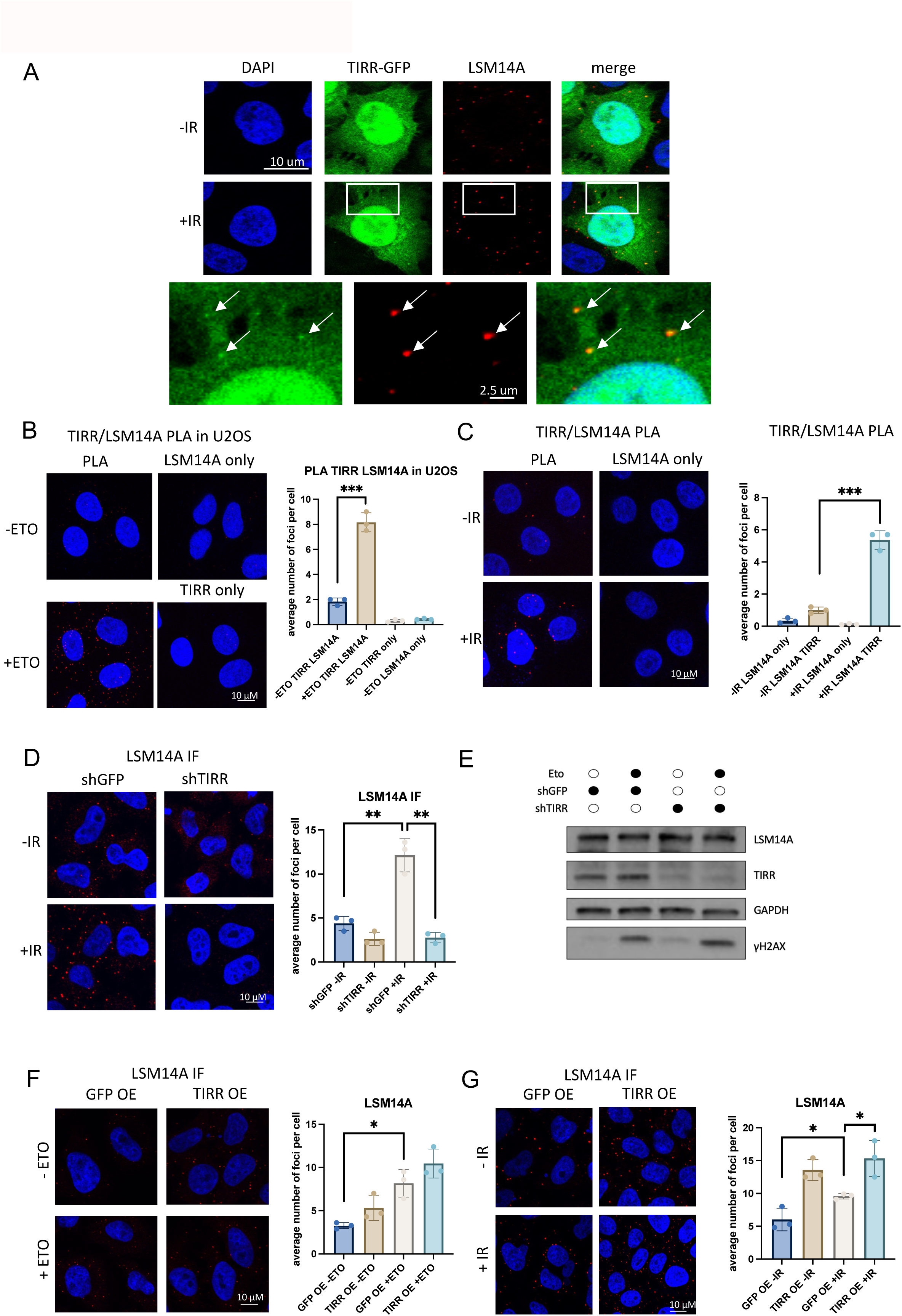

**Supplementary Figure 11.**
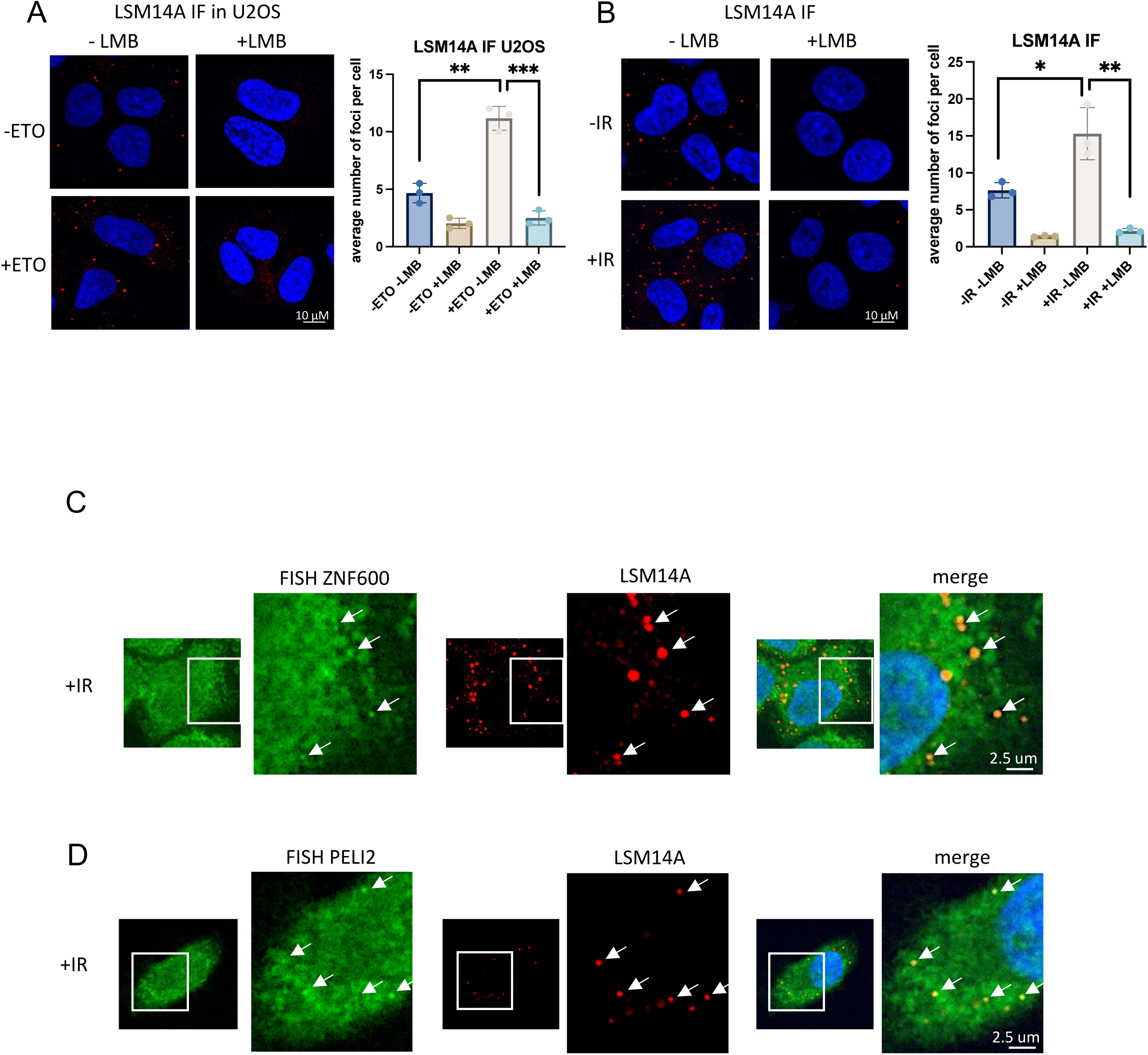

**Supplementary Figure 12.**
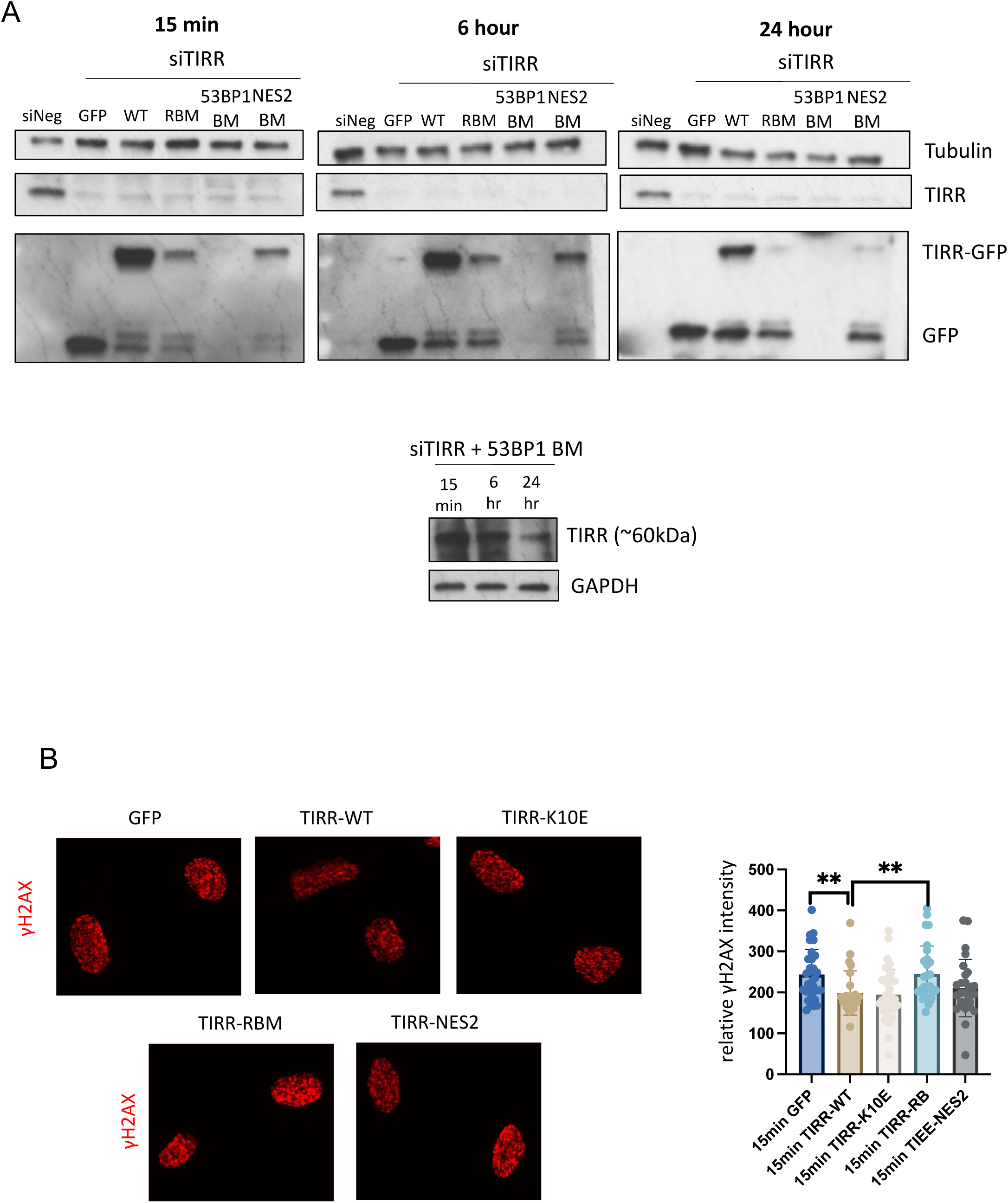

