## Supplementary Figure legends for "TIRR regulates mRNA export and association with P bodies in response to DNA damage"

**Supplementary Figure 1**

A) Overview of UV RIP-Seq protocol. Made with Biorender.com.

B) TIRR-GFP Flp-IN Trex cells were induced to express TIRR-GFP using Doxycycline at different time points up to 24 hours. A time point of 16 hours was used for RIP-Seq.

C) Analysis of read counts in RIP-seq samples from TIRR-GFP or GFP pull downs from cells with or without Etoposide (ETO) treatment.

D) Heat map analysis showing similarity between RIP-seq samples.

E) Principal Component analysis of top and bottom differentially expressed genes.

F) Principal Component analysis (PCA)of RIP-seq samples.

G) Scree plot analysis of RIP-seq samples.

**Supplementary Figure 2**

A) GO analysis of mRNA bound to TIRR in no damage condition.

B) String analysis of mRNA bound to TIRR in damage (ETO) condition.

C) Distribution of transcription factors genes bound to TIRR in damage condition across human genome.

**Supplementary Figure 3**

Comparison of features of RNAs bound to TIRR in damage conditions vs the reference genome. Features include: coding sequence length, 5'UTR length, transcript length, 3' UTR length and genome span.

**Supplementary Figure 4**

A) Heatmap analysis showing similarity between RNA-seq samples from cells expressing shGFP (control) or shTIRR, with or without Etoposide (ETO) treatment.

B) Principal component analysis (PCA) of RNA-seq samples, as in A).

C) Dot plot showing expression levels of TIRR in RNA-seq samples, as in A).

D) Volcano plot showing differentially expressed RNA in samples from cells expressing shTIRR and shGFP in no damage condition.

E) Volcano plot showing differentially expressed RNA in samples from cells expressing shTIRR and shGFP in damage (ETO) condition.

F) Volcano plot showing differentially expressed RNA in samples from cells expressing shGFP in no damage (no ETO) and damage (ETO) condition.

G) Volcano plot showing differentially expressed RNA in samples from cells expressing shTIRR in no damage (no ETO) and damage (ETO) condition.

**Supplementary Figure 5**

A) Principal component analysis (PCA) of RNA-seq samples from cells expressing shGFP (control) or shTIRR, with or without IR treatment. DNA damage was induced by IR and samples were collected 1 hour later.

B) Dot plot showing expression levels of TIRR in RNA-seq samples, as in A).

C) Volcano plot showing differentially expressed RNA in samples from cells expressing shTIRR and shGFP in no damage (no IR) condition.

D) Volcano plot showing differentially expressed RNA in samples from cells expressing shTIRR and shGFP in damage (IR) condition.

E) Volcano plot showing differentially expressed RNA in samples from cells expressing shGFP in no damage (no IR) and damage (IR) condition.

F) Volcano plot showing differentially expressed RNA in samples from cells expressing shTIRR in no damage (no IR) and damage (IR) condition.

G) Principal component analysis (PCA) of RNA-seq samples from cells expressing shGFP (control) or shTIRR, with or without damage treatment comparing ETO and IR.

**Supplementary Figure 6**

A) Representative western blot showing protein levels of IRF3, HOXD11, TIRR, γH2AX and Tubulin in cells expressing shGFP and shTIRR with or without Etoposide treatment.

B) Quantification of HOXD11 protein levels, n=3, as in A)

C) Quantification of IRF3 protein levels, n=3, as in A)

**Supplementary Figure 7**

A) Left: Confocal images showing RNA FISH signals (in green) for *ZNF600* in shGFP or shTIRR cells in no damage (-ETO) condition. Right: Quantification of *ZNF600* mRNA signals in nucleus and cytoplasm plotted as relative FISH signal intensity in shGFP or shTIRR cells upon ETO, n>50 cells. Significance was determined using t- test, (***$p \leq 0.001$, ****$p \leq 0.0001$).

B) As in A) for *PELI2* mRNA.
C) As in A) for *SPEN* mRNA
D) As in A) for *Tubulin* mRNA

**Supplementary Figure 8**

A) Left: Confocal images showing RNA FISH signals (in green) for *ZNF600* in shGFP or shTIRR cells in damage (+IR) condition. Right: Quantification of *ZNF600* mRNA signals in nucleus and cytoplasm plotted as relative FISH signal intensity in shGFP or shTIRR cells upon IR, n>50 cells. Significance was determined using t- test, (***$p \leq 0.001$, ****$p \leq 0.0001$).
B) As in A) for *PELI2* mRNA.
C) Western blot of Co-IP of XPO1 with TIRR-GFP in damage and non-damage conditions. A GFP only control was used to assess non-specific binding of XPO1 to the beads.
D) Left: Confocal images showing PLA of TIRR and XPO1 in cells with or without IR treatment. Single antibody was used as negative control. Right: Quantification of PLA signals. Significance was determined using non-parametric Mann-Whitney test (****$p \leq 0.0001$).
E) Confocal images showing PLA of TIRR and XPO1 in cells expressing GFP control. Single antibodies were used as negative controls.

**Supplementary Figure 9**

A) Left: Representative confocal images showing immunofluorescence of TIRR in HeLa cells with or without IR treatment and with or without LMB treatment. DAPI was used to stain nuclei. Right: Quantification of TIRR signal in cytoplasm. n>50 cells, significance was determined using t-test, (****$p \leq 0.0001$).
B) Left: Representative confocal images showing immunofluorescence of TIRR in U2OS cells with or without Etoposide (ETO) treatment and with or without LMB treatment. DAPI was used to stain nuclei. Right: Quantification of TIRR signal in cytoplasm. n>50 cells, significance was determined using t-test, (****$p \leq 0.0001$).
C) Western blot showing expression levels of TIRR-GFP, TIRR, mCherry-LaminB1-10, LaminB1 and GAPDH in U2OS cells transfected with PA-GFP-TIRR and mCherry-LaminB1-10.

**Supplementary Figure 10**

A) Representative confocal images showing immunofluorescence of TIRR in cells with or without IR treatment. TIRR expression was visualised in green channel (GFP), P-bodies were labelled using anti-LSM14A antibody and visualised in red channel. White arrows point to P-bodies.

B) Left: proximity ligation assay using anti-TIRR and anti-LSM14A antibodies in no damage (-ETO) or damage (+ETO) conditions in U2OS cells. DAPI was used to label nuclei. PLA signals were visualised in red channel. Single antibodies were used as negative controls. Right: quantification of top. Error bar = mean ±SD, significance was determined using non-parametric Mann Whitney test (***$p \leq 0.001$).

C) Left: proximity ligation assay using anti-TIRR and anti-LSM14A antibodies in no damage (-IR) or damage (+IR) conditions in HeLa cells. DAPI was used to label nuclei. PLA signals were visualised in red channel. Single antibodies were used as negative controls. Right: quantification of top. Error bar = mean ±SD, significance was determined using non-parametric Mann Whitney test, (***$p \leq 0.001$).

D) Left: Immunofluorescence analysis showing LSM14A staining in no damage (-IR), and damage (+IR) conditions in cells expressing either shGFP (control) or shRNA targeting TIRR (shTIRR). DAPI was used to stain nuclei. Right: Quantification of TIRR signal in cytoplasm, n>50 cells, significance was determined using t- test, (**$p \leq 0.01$).

E) Western blot showing protein levels of LSM14A, TIRR, GAPDH and γH2AX in cells expressing either shGFP (control) or shRNA targeting TIRR (shTIRR).

F) Left: Immunofluorescence analysis showing LSM14A staining in no damage (-ETO), and damage (+ETO) conditions in cells over-expressing either GFP (control) or TIRR. DAPI was used to stain nuclei. Right: Quantification of TIRR signal in cytoplasm, n>50 cells, significance was determined using t- test, (*$p \leq 0.05$).

G) Left: Immunofluorescence analysis showing LSM14A staining in no damage (-IR), and damage (+IR) conditions in cells over-expressing either GFP (control) or TIRR. DAPI was used to stain nuclei. Right: Quantification of TIRR signal in cytoplasm, n>50 cells, significance was determined using t- test, (*$p \leq 0.05$).

**Supplementary Figure 11**

A) Left: Immunofluorescence analysis showing LSM14A staining in no damage (-ETO), and damage (+ETO) conditions in U2OS cells with or without LMB treatment. DAPI was used to stain nuclei. Right: Quantification of TIRR signal in cytoplasm, n>50 cells, significance was determined using t- test, (**$p \leq 0.01$, ***$p \leq 0.001$).

B) Left: Immunofluorescence analysis showing LSM14A staining in no damage (-IR), and damage (+IR) conditions in HeLa cells with or without LMB treatment. DAPI was used to stain nuclei. Right: Quantification of TIRR signal in cytoplasm, n>50 cells, significance was determined using t- test, (*$p \leq 0.05$, **$p \leq 0.01$).

C) Representative confocal images showing co-localisation of RNA FISH signals (in green) for *ZNF600* with LSM14A (in red) in HeLa cells after IR treatment. n>50 cells. White rectangles mark zoomed in images on the right. White arrows point to P-bodies with FISH signal (in yellow).

D) As in A) for *PELI2* mRNA.

**Supplementary Figure 12**

A) Western blot showing levels of (top panels) endogenous TIRR and TIRR-GFP expressed from plasmids (right) in cells 15min, 6 hours and 24 hours post IR treatment. Tubulin was used as a loading control. Bottom panels showing expression of TIRR-K10E-NeonGreen (53BP1 BM).

B) Immunofluorescence showing γH2AX signal in cells transfected with GFP, TIRR-WT, TIRR-K10E, TIRR-RBM and TIRR-NES2 mutant, at 15min post IR treatment. Significance was determined using t- test, (**$p \leq 0.01$, ****$p \leq 0.0001$).
