## Supplementary Table 3 for "TIRR regulates mRNA export and association with P bodies in response to DNA damage"

**Sequences**

| **Oligo** | **Sequence** | **Use** | **Modification** |
| --- | --- | --- | --- |
| PELI2 FISH PROBE | AAGCACCATTGTACCCGAGC | FISH | 5' Alexa Fluor NHS ester |
| ZNF600 FISH PROBE | TGCGTTTGGAAGAGATATCCAC | FISH | 5' Alexa Fluor NHS ester |
| SPEN FISH PROBE | AGCGCTCCAAACGAGACTTG | FISH | 5' Alexa Fluor NHS ester |
| BETA-TUBULIN FISH PROBE | CAGAGTCCATGGTCCCAGGT | FISH | 5' Alexa Fluor NHS ester |
| TIRR_BACK_FWD | TGGACGAGCTGTACAAGTAACAATTCACTCGATCGGCTCGCTGATC | Gibson Cloning PAGFP-TIRR (TIRR backbone) | None |
| TIRR_BACK_REV | AGCTCCTCGCCCTTGCTCACCTCGACATCGATTGCGGCCGCAGAG | Gibson Cloning PAGFP-TIRR (TIRR backbone) | None |
| PA-GFP_FWD | CGGCCGCAATCGATGTCGAGGTGAGCAAGGGCGAGGAGCTGTTCA | Gibson Cloning PAGFP-TIRR (PA-GFP fragment) | None |
| PA-GFP_REV | CGAGCCGATCGAGTGAATTGTTACTTGTACAGCTCGTCCATGCCG | Gibson Cloning PAGFP-TIRR (PA-GFP fragment) | None |
| MSCARLET_FWD | CGCTACCGGTCGCCACCATGGTGAGCAAGGGCGAGGCAGTGATCAAG | Gibson Cloning LSM14A-mScarlet (mScarlet fragment) | None |
| MSCARLET_REV | TTGAGCTCGAGATCTGAGTACTTGTACAGCTCGTCCATGCCGCCG | Gibson Cloning LSM14A-mScarlet (mScarlet fragment) | None |
| LSM14A_BACK_FWD | GCATGGACGAGCTGTACAAGTACTCAGATCTCGAGCTCAAGCTTCGAATTCCCATG | Gibson Cloning LSM14A-mScarlet (LSM14A backbone) | None |
| LSM14A_BACK_REV | ACTGCCTCGCCCTTGCTCACCATGGTGGCGACCGGTAGCGCTAGC | Gibson Cloning LSM14A-mScarlet (LSM14A backbone) | None |
| FRAG_MRB_FWD | CGGGCTGGAGCCACTCGTGCGCCGCCATGCTGTACGCCGCCAACCCTGGGCAGCTCTTCGGCCGCATCCCCATGCGCTTCTCGGTGCTGATGCAGATGGCCTTCGACGGGCTGCTGGGCTTCCCCGGGGGCGCCGTGGACCGGCGCTTCTGGTCGCTGGAGGACGGCCTGAACCGGGTGCTGGGCCTGGGCCTGGGCTGC | Generating fragment for RBM TIRR | None |
| FRAG_MRB_REV | ACCCGCACGAGGCCCAGCACGGCCAGGCCGTGGTCGCGCGAGTGCACCGCGCTGATCTCCACGGCGTGCAGCTGCTCCAGCGTCAGCTGCCGCGCGTACAGGTGCGCCACGACGCGGTGTGGGCCCTCGGTCAGGTGCGAGCTCAGGTAGTCGGCCTCGGTGAGGCGCAGGCAGCCCAGGCCCAGGCCCAGCACCCGGTT | Generating fragment for RBM TIRR | None |
| BACK_EGFP_TIRR_MRB_FWD | GTGCTGGGCCTCGTGCGGGTCCCGCTGTACACCCAGAAGGAC | Gibson Cloning LSM14A-mScarlet (GFP backbone) | None |
| BACK_EGFP_TIRR_MRB_REV | GCACGAGTGGCTCCAGCCCGGCCCTAGGCGCATCGCCTCC | Gibson Cloning LSM14A-mScarlet (GFP backbone) | None |
| MRB_TIRR_FWD | GGAGGCGATGCGCCTAGGGCCGGGCTGGAGCCACTCGTGC | Gibson Cloning LSM14A-mScarlet (MRB TIRR fragment) | None |
| MRB_TIRR_REV | CCTTCTGGGTGTACAGCGGGACCCGCACGAGGCCCAGCAC | Gibson Cloning LSM14A-mScarlet (MRB TIRR fragment) | None |
| TIRR_WT_RED_FWD | AGGTCTATATAAGCAGAGCTCAAGCTTGCGGCCGCCACCATGTCGACGGCGGCGGTTCCGGAGCTGAAGCAGATCAGCCGGGTGGAGGCGATGCGCCTAGGGCCGGGCTGGAGCCACTCGTGCCACGCCATGCTGTACGCCGCCAACCCTGGGCAGCTCTTCGGCCGCATCCCCATGCGC | Generating fragment for green NES mutant (WT segment) | None |
| TIRR_MUT_RED_FWD | AGGTCTATATAAGCAGAGCTCAAGCTTGCGGCCGCCACCATGTCGACGGCGGCGGTTCCGGAGGCCAAGCAGGCCAGCCGGGCCGAGGCGGCCCGCGCCGGGCCGGGCTGGAGCCACTCGTGCCACGCCATGCTGTACGCCGCCAACCCTGGGCAGCTCTTCGGCCGCATCCCCATGCGC | Generating fragment for red NES mutant (mutant segment) | None |
| TIRR_WT_GREEN_REV | CAGGTAGTCGGCCTCGGTGAGGCGCAGGCAGCCCAGGCCCAGGCCCAGCACCCGGTTCAGGCCGTCCTCCAGCGACCAGAAGCGCCGGTCCACGAAGCCCCCGGGGAAGCCCAGCAGCCCGTCGAAACGCATCTGCATCAGCACCGAGAAGCGCATGGGGATGCGGCCGAAGAGCTGCCC | Generating fragment for red NES mutant (WT segment) | None |
| TIRR_MUT_GREEN_REV | CAGGTAGTCGGCCTCGGTGAGGCGCAGGCAGCCGGCGCCGGCGCCGGCGGCCCGGTTGGCGCCGTCCTCGGCCGACCAGAAGCGCCGGTCCACGAAGCCCCCGGGGAAGCCCAGCAGCCCGTCGAAACGCATCTGCATCAGCACCGAGAAGCGCATGGGGATGCGGCCGAAGAGCTGCCC | Generating fragment for green NES mutant (mutant segment) | None |
| TIRR_WT_BLUE_FWD | CTGAGCTCGCACCTGACCGAGGGCCCACACCGCGTCGTGGCGCACCTGTACGCGCGGCAGCTGACGCTGGAGCAGCTGCACGCCGTGGAGATCAGCGCGGTGCACTCGCGCGACCACGGCCTGGAGGTGCTGGGCCTCGTGCGGGTCCCGCTGTACACCCAGAAGGACCGAGTCGGAGGC | Generating fragment for orange NES mutant (WT segment) | None |
| TIRR_MUT_BLUE_FWD | CTGAGCTCGCACCTGACCGAGGGCCCACACCGCGCCGCCGCGCACGCCTACGCGCGGCAGGCCACGGCCGAGCAGCTGCACGCCGTGGAGATCAGCGCGGTGCACTCGCGCGACCACGGCCTGGAGGTGCTGGGCCTCGTGCGGGTCCCGCTGTACACCCAGAAGGACCGAGTCGGAGGC | Generating fragment for blue NES mutant (mutant segment) | None |
| TIRR_WT_ORANGE_REV | GGTGGCTGCAGCCAGGGCCTCAACCAGCTTCTCCTCGGGCATCATGTTGAGCACCTTGAGGGCAAAGAGGAGCTGGCACTTAGCCGTGCTCACGAAGGCGTTGCTCAGGAAGTTGGGGAAGCCTCCGACTCGGTCCTTCTGGGTGTACAGCGGGACCCGCACGAGGCCCAGCACCTCCAG | Generating fragment for blue NES mutant (WT segment) | None |
| TIRR_MUT_ORANGE_REV | GGTGGCTGCAGCCAGGGCCTCAACCAGCTTCTCCTCGGGGGCGGCGTTGGCCACCTTGGCGGCAAAGGCGGCCTGGCACTTAGCCGTGCTCACGAAGGCGTTGCTCAGGAAGTTGGGGAAGCCTCCGACTCGGTCCTTCTGGGTGTACAGCGGGACCCGCACGAGGCCCAGCACCTCCAG | Generating fragment for orange NES mutant (mutant segment) | None |
| NES_RG_FWD | TGCCTGCGCCTCACCGAGGCC | Gibson Cloning red and green NES mutant | None |
| NES_RG_REV | TGGTGGCGGCCGCAAGCTTGA | Gibson Cloning red and green NES mutant | None |
| NES_BO_FWD | AAGCTGGTTGAGGCCCTGGCTGC | Gibson Cloning blue and orange NES mutant | None |
| NES_BO_REV | GGCCCTCGGTCAGGTGCGAGC | Gibson Cloning red and green NES mutant | None |
